# Task-dependent representations of stimulus and choice in mouse parietal cortex

**DOI:** 10.1101/144592

**Authors:** Gerald N. Pho, Michael J. Goard, Jonathan Woodson, Benjamin Crawford, Mriganka Sur

## Abstract

The posterior parietal cortex (PPC) has been implicated in perceptual decisions, but whether its role is specific to sensory processing or sensorimotor transformation is not well understood. To distinguish these possibilities, we trained mice of either sex to perform a visual discrimination task and imaged the activity of PPC populations during both engaged behavior and passive viewing. Unlike neurons in primary visual cortex (V1), which responded robustly to stimuli in both conditions, most neurons in PPC responded exclusively during task engagement. However, PPC responses were heterogeneous, with a smaller subset of neurons exhibiting stimulus-driven, contrast-dependent responses in both conditions. Neurons in PPC also exhibit stronger modulation by noise correlations relative to V1, as illustrated by a generalized linear model that takes into account both task variables and between-neuron correlations. To test whether PPC responses primarily encoded the stimulus or the learned sensorimotor contingency, we imaged the same neurons before and after re-training mice on a reversed task contingency. Unlike V1 neurons, most PPC neurons exhibited a dramatic shift in selectivity after re-training and reflected the new sensorimotor contingency, while a smaller subset of neurons preserved their stimulus selectivity. Mouse PPC is therefore strongly task-dependent, contains heterogeneous populations sensitive to stimulus and choice, and may play an important role in the flexible transformation of sensory inputs into motor commands.

**Significance Statement:** Perceptual decision making involves both processing of sensory information and mapping that information onto appropriate motor commands via learned sensorimotor associations. While visual cortex (V1) is known to be critical for sensory processing, it is unclear what circuits are involved in the process of sensorimotor transformation. While the mouse posterior parietal cortex (PPC) has been implicated in visual decisions, its specific role has been controversial. By imaging population activity while manipulating task engagement and sensorimotor contingencies, we demonstrate that PPC, unlike V1, is highly task-dependent, heterogeneous, and sensitive to the learned task demands. Our results suggest that PPC is more than a visual area, and may instead be involved in the flexible mapping of visual information onto appropriate motor actions.

## Introduction

Perceptual decision-making involves multiple cognitive processes, including processing of sensory stimuli, accumulation of evidence, and transformation of sensory information into an appropriate motor plan. Although many brain regions have been implicated in perceptual decisions, dissociating their individual contribution to these different processes remains a challenge.

In particular, the posterior parietal cortex (PPC) has been hypothesized to play a key role in at least some types of decision tasks in both primates (Shadlen and Newsome, 2001; Gold and Shadlen, 2007) and in rodents (Harvey et al., 2012; Raposo et al., 2014; Goard et al., 2016). However, the specific role of rodent parietal cortex, and whether its function is homologous to that of primates, remains unclear. Recent electrophysiological recordings in rats during auditory decisions (Hanks et al., 2015) recapitulated findings in primates that PPC neurons can reflect the accumulated evidence for a decision. However, pharmacological inactivation of PPC during the same auditory task has no effect on behavioral performance (Erlich et al., 2015), and a minimal role of PPC was also found for whisker-based decisions in mice (Guo et al., 2014). These findings are in contrast to studies involving visual decision tasks, where activity in PPC not only encodes information about the decision (Raposo et al., 2014; Goard et al., 2016; Morcos and Harvey, 2016) but is also causally necessary for behavior (Harvey et al., 2012; Goard et al., 2016; Licata et al., 2017).

The apparent specificity of behavioral deficits to the visual modality has led some to argue that rodent PPC may ultimately be more homologous to extrastriate cortex in processing sensory signals that are accumulated elsewhere for decision-making (Licata et al., 2017). Indeed, both anatomical projection studies (Wang and Burkhalter, 2007; Wang et al., 2012), as well as functional mapping studies (Marshel et al., 2011; Garrett et al., 2014) indicate that PPC may overlap with or contain a group of retinotopically-organized extrastriate areas that are rostral to V1. An alternative possibility is that PPC may play a specific role in the mapping of visual stimuli to motor commands. If this were the case, one may expect that activity in PPC would be highly task-dependent, and may even flexibly re-map depending on learned sensorimotor contingencies (Freedman and Assad, 2006).

To distinguish these possibilities, we used population calcium imaging to measure activity in PPC and in the primary visual cortex (V1) during a go/no-go lick-based visual discrimination task. Having previously demonstrated the necessity of PPC in the performance of this task (Goard et al., 2016), we sought in this work to investigate its specific role in either sensory processing or sensorimotor transformation by manipulating task engagement and learned task contingencies. V1 neurons had robust visual responses during both task engagement and passive viewing of stimuli that remained stable after task contingency reversal. By contrast, PPC responses were largely specific to task performance and were highly sensitive to task contingency. Our results are consistent with a role of the mouse posterior parietal cortex in transforming sensory information to motor commands during perceptual decisions.

## Materials and Methods

### Mice and surgery

All experiments were carried out in mice of either sex using protocols approved by Massachusetts Institute of Technology’s Animal Care and Use Committee and conformed to National Institutes of Health guidelines. Data were collected from adult (3-5 months old) wild-type (C57BL/6; n = 15) mice of either sex. The animals were housed on a 12 hour light/dark cycle in cages of up to 5 animals before the implants, and individually after the implants. All surgeries were conducted under isoflurane anesthesia (3.5% induction, 1.5-2.5% maintenance). Meloxicam (1 mg kg^-1^, subcutaneous) was administered pre-operatively and every 24 hours for 3 days to reduce inflammation. Once anesthetized, the scalp overlying the dorsal skull was sanitized and removed. The periosteum was removed with a scalpel and the skull was abraded with a drill burr to improve adhesion of dental acrylic. Stereotaxic coordinates for future viral injections were marked with a non-toxic ink and covered with a layer of silicon elastomer (Kwik-Sil, World Precision Instruments) to prevent acrylic bonding. The entire skull surface was then covered with dental acrylic (C&B-Metabond, Parkell) mixed with black ink to reduce light transmission. A custom-designed stainless steel head plate (eMachineShop.com) was then affixed using dental acrylic. After head plate implantation, mice recovered for at least five days before beginning water restriction.

After behavioral training was complete, animals were taken off water restriction for five days before undergoing a second surgery to implant the imaging window. Procedures for anesthetic administration and post-operative care were identical to the first surgery. The dental acrylic and silicon elastomer covering the targeted region were removed using a drill burr. The skull surface was then cleaned and a craniotomy was performed over left V1/PPC, leaving the dura intact. Neurons were labeled with a genetically-encoded calcium indicator by microinjection (Stoelting) of 50 nl AAV2/1.Syn.GCaMP6s.WPRE.SV40 (University of Pennsylvania Vector Core; diluted to a titer of 10^12^ genomes ml^-1^) 300 μm below the pial surface. Between two and five injections were made in each exposed region, centered at V1 (4.2 mm posterior, 2.5 mm lateral to Bregma) and PPC (2 mm posterior, 1.7 mm lateral to Bregma). Since the viral expression spreads laterally from the injection site, exact stereotaxic locations were photographed through the surgical microscope for determining imaging areas. Finally, a cranial window was implanted over the craniotomy and sealed first with silicon elastomer then with dental acrylic. The cranial windows were made of two rounded pieces of coverglass (Warner Instruments) bonded with optical glue (NOA 61, Norland). The bottom piece was a circular coverglass (4 mm diameter) that fit snugly in the craniotomy. The top piece was a larger circular coverglass (3-5 mm, depending on size of bottom piece) and was bonded to the skull using dental acrylic. Mice recovered for five days before commencing water restriction.

### Behavioral tasks

We trained mice to perform a head-fixed go/no-go visual discrimination task, similar to previous designs (Goard et al., 2016). Stimuli consisted of full-contrast sine wave gratings (spatial frequency: 0.05 cycles deg^-1^; temporal frequency: 2 Hz) drifting at either 0° (upwards, target, Stimulus A) or 90° (rightwards, non-target, Stimulus B) away from vertical. Stimuli were presented to the right eye alone by placing the screen at an oblique angle to the animal. Behavioral training and testing was implemented with custom software written in Matlab (Mathworks) using Psychtoolbox-3 (Kleiner et al., 2007) and Data Acquisition toolbox. Spout position was controlled by mounting the spout apparatus on a pneumatically-driven sliding linear actuator (Festo) controlled by two solenoids. Licks were detected using an infrared emitter/receiver pair (Digikey) mounted on either side of the retractable lick spout. Mice were water-restricted and earned most of their daily ration (1mL) during training.

An auditory cue tone (5 kHz, 0.5 s, 65 dB SPL) indicated the beginning of each trial. After a 1 s delay, a visual stimulus was presented for 2 s. At the end of the stimulus epoch, the spout was rapidly moved within reach of the tongue, and remained within reach for 1.5 s. Correct licks during this period were rewarded with 5-8 μl water and a brief reward tone (10 kHz, 0.1 s). Licks to the non-target stimulus were punished with a white noise auditory stimulus alone (early training) or white noise plus 1-3 μl of 5mM quinine hydrochloride in water (late training). This concentration was chosen to deter licking to non-targets without causing mice to lose motivation altogether. At the end of the response epoch, the spout was then rapidly retracted and remained out of reach until the next trial (3 s inter-trial interval).

Mice were trained in successive stages, as previously described (Goard et al., 2016). Once mice reached high levels of performance at the final stage of the task (*d’ >* 1.5 and *R_HIT_ - R_FA_* > 50%), they were removed from water restriction for window implantation. Mice reached criterion performance after an average of 92 ± 11 sessions. After recovery from window implantation surgery, they were re-trained to a level of high performance (2-7 days) before beginning experimental sessions. Any sessions with poor performance were discarded (minimum performance criterion: *d’* > 1 and *R_HIT_ - R_FA_* > 30%).

During imaging experiments, blocks of engaged behavior trials were alternated with blocks of passive viewing. Blocks were 5-10 min in duration (40-80 trials per block). During passive blocks, the spout was out of reach for the duration of the block. A few extra passive trials were given (without imaging) before each passive block to ensure that mice did not expect spout presentation during all imaged passive trials. The sequence of target and non-target stimuli presented for a given passive block was matched to the sequence of stimuli used for the preceding engaged block. In some cases, instead of alternating between engaged and passive blocks, the engaged blocks were all grouped together at the beginning of the session, followed by an equal number of consecutive passive blocks. No difference in results was observed for alternating versus grouped blocks.

Some mice (n = 6) were trained on a variable contrast version of the task. On each trial, the stimulus was randomly set to one of six contrasts (2, 4, 8, 16, 32, or 64%), regardless of whether the stimulus was a target or non-target. The mouse therefore could not predict the contrast of the stimulus from trial to trial.

Some mice (n = 3) were re-trained after initial imaging experiments on a reversed reward contingency. Reward contingency was switched abruptly, with reward given for licks to Stimulus B and no reward for licks to Stimulus A. Because mice were quickly discouraged by the reversal, no quinine punishment was initially given. Additionally, the reward tone (10 kHz, 0.1 s) was paired with the onset of the new target stimulus (Stimulus B) early in re-training, in order to encourage licking. Three of the five mice trained on this reversed contingency achieved criterion performance after re-training for 10 ± 2 days; the other two were removed from the study.

For all tasks, behavioral d-prime was computed by norminv(Hit rate) – norminv(False alarm rate), where norminv() is the inverse of the cumulative normal function (Green and Swets, 1966; Carandini and Churchland, 2013). Values of Hit and False alarm rate were truncated between 0.01 and 0.99, setting the maximum d’ to 4.65. For illustration purposes, d-prime before and after reversal was computed using Stimulus A as the target in both conditions (**Figure 9B**).

### Two-photon imaging

GCaMP6s fluorescence was imaged 14-35 days after virus injection using Prairie Ultima IV 2-photon microscopy system with a resonant galvo scanning module (Bruker). For fluorescence excitation, we used a Ti-Sapphire laser (Mai-Tai eHP, Newport) with dispersion compensation (Deep See, Newport) tuned to *λ* = 910 nm. For collection, we used GaAsP photomultiplier tubes (Hamamatsu). To achieve a wide field of view, we used a 16x/0.8 NA microscope objective (Nikon), which was mounted on a Z-piezo (Bruker) for volume scanning. An optical zoom of 2x was used in most cases to improve spatial resolution. Resonant scanning (15.9 kHz line rate, bidirectional) was synchronized to z-piezo steps in the acquisition software for volume scanning. For volume scanning, four 441 x 512 pixel imaging planes separated by 20 or 25 μm were imaged sequentially at a stack rate of 5 Hz in 5-10 min imaging blocks. There was very little redundant sampling of neurons between imaging planes (<1%) as assayed by correlation coefficient of spontaneous activity. Laser power ranged from 40-75 mW at the sample depending on GCaMP6s expression levels. Photobleaching was minimal (<1% min^-1^) for all laser powers used. A custom stainless steel plate (eMachineShop.com) attached to a black curtain was mounted to the head plate before imaging to prevent light from the visual stimulus monitor from reaching the PMTs. During imaging experiments, the polypropylene tube supporting the mouse was suspended from the behavior platform with high tension springs (Small Parts) to dampen movement.

### Image preprocessing and cell selection

Calcium imaging data were acquired using PrairieView acquisition software and sorted into multipage TIF files. All analyses were performed using custom scripts written either in ImageJ or MATLAB (Mathworks).

Images were first corrected for X-Y movement by registration to a reference image (the pixel-wise mean of all frames) using 2-dimensional cross correlation. To identify responsive neural somata, a pixel-wise activity map was calculated as previously described (Ahrens et al., 2012). Neuron cell bodies were identified using local adaptive threshold and iterative segmentation. Automatically-defined ROIs were then manually checked for proper segmentation in a MATLAB-based graphical user interface (allowing comparison to raw fluorescence and activity map images). To subtract the influence of local neuropil on somatic signals, the fluorescence in the somata was estimated as *F_corrected_soma_(t) = F_raw_soma_ (t) - 0.7 × F_neuropii_(t)*, where *F_neuropii_* was the defined as the fluorescence in the region 0-15 mm from the ROI border (excluding other ROIs) (Chen et al., 2013). *ΔF/F* for each neuron was calculated as *ΔF/F_t_ = (F_t_-F_0_)/F_0_,* with *F_0_* defined as the mode of the raw fluorescence density distribution.

To align ROIs between different imaging sessions across days (**Figure 9, Figure 10**), we used a semi-automated method similar to prior work (Huber et al., 2012). First, for each plane, anchor points were manually defined by visual comparison of the two average projection images. These anchor points helped to define a predicted displacement vector field that would be used to map coordinates from one session to the other. For each coordinate, the predicted vector was defined by the average (weighted inversely by distance) of the vectors for all defined anchor points.

Next, for each ROI, a square region (~4x the size of the ROI) around the ROI was selected. To determine the displacement across sessions, we computed the normalized cross-correlation of this square with the average projection of the other session. This was multiplied point-by-point with a mask that decayed gradually with distance from the predicted displacement vector, and then smoothed with a 2-D Gaussian filter. The peak of the resulting image was taken to be the actual displacement vector of the ROI. This process biases the displacement of each ROI towards the vector predicted from the manually-defined anchor points. Finally, any ROIs with a computed displacement vector that differed by greater than 5 pixels from the predicted vector were flagged for manual inspection, and then either redrawn or removed.

After image preprocessing and *ΔF/F* extraction, traces were sorted by trial type (hit, miss, correct reject, false alarm) and condition (engaged, passive). The baseline response (1 s before stimulus onset) was subtracted from each trial. A neuron was considered task responsive if its mean *ΔF/F* during the last 1.6 s (8 frames) of the stimulus period was significantly (p < 0.01, t-test) greater than the pre-stimulus baseline (1 s), for either hit or correct reject trials. Neurons also had to meet a signal-to-noise criterion, needing a trial-averaged response that exceeded a threshold of at least two standard deviations above baseline during either the stimulus or choice period. All further analyses are based on responses during the last 1.6 s of the stimulus period, unless noted otherwise. Cell selection criteria for error analyses, variable contrast analyses, and reversal analyses are described in the appropriate sections below.

### Experimental design and statistical analysis

Data was obtained from 15 mice, 5 with V1 only, 6 with PPC only, and 4 with both V1 and PPC. For most mice, multiple fields of view were sampled within V1 or within PPC. Each field was imaged for a single session, consisting of multiple Engaged and Passive blocks and yielding on average 94 trials (minimum 47) per stimulus per condition. For variable contrast tasks, an average of 25 trials (minimum 8) were acquired per contrast. Significantly task-responsive cells from different fields were pooled by area. No tests were conducted to determine sample size. For the full-contrast task, data came from 1915 V1 cells (18 fields, 9 mice) and 3524 PPC cells (22 fields, 10 mice). For the variable-contrast task, data came from 250 V1 cells (8 fields, 3 mice) and 611 PPC cells (11 fields, 6 mice). For comparisons before and after contingency reversal, data came from 488 V1 cells (8 fields, 3 mice) and 509 PPC cells (8 fields, 3 mice).

All statistical analysis was performed using custom-written scripts in MATLAB or R. In all cases, data was not assumed to be normal, and nonparametric and/or permutation tests (2000 permutations) were used to assess statistical significance of results. All tests were two-tailed, and a significance level of p < 0.05 was considered significant. Unless otherwise noted, all measures are reported as mean ± SEM. When estimating the percentage of selective or modulated neurons, bootstrapping across imaging fields was used to generate confidence intervals on the percentages for each area. When testing the statistical significance of differences between V1 and PPC neurons that were pooled across imaging fields, clustered nonparametric tests (Datta and Satten, 2005, 2008) were used to account for intra-cluster correlations (Galbraith et al., 2010; Aarts et al., 2014) using the *clusrank* package in R.

### Selectivity and modulation indices

Neurons were marked as target- or non-target-preferring (**Figure 2E**) based on their mean response during Engaged trials. Neurons were marked as task-gated if they did not exhibit a significant response to their preferred stimulus during Passive trials (**Figure 2F**).

All comparative indices (engagement modulation index, error modulation index, contrast modulation index, selectivity index) were computed using a receiver operating characteristic (ROC) analysis, which quantifies the ability of an ideal observer to discriminate between trial types based on single trial responses (Green and Swets, 1966; Britten et al., 1996). Each index was derived from the area under the ROC curve (auROC), and defined as 2 × (auROC - 0.5); this value ranged from -1 to 1 (Raposo et al., 2014). Engagement modulation index (**Figure 2H**, **Figure 6E**, **F**, **Figure 10D**) was computed by comparing the stimulus period response (last 1.6 s) during engaged and passive conditions. A value of +1 indicates that a cell always fired more on Engaged trials. Error modulation index (**Figure 3C**) was computed by comparing False Alarm trials to Correct Reject trials, with positive values indicating stronger response on FA trials. Contrast modulation index (**Figure 6C**) was computed by comparing responses at the highest and lowest contrasts, with positive values indicating preference for high contrasts. Selectivity index (**Figure 2G**) was computed by comparing target and non-target responses, with positive values indicating preference for target stimuli. Finally, selectivity index was computed separately before and after reversal training (**Figure 9E-H**) by comparing responses to the Stimulus A (original target) and Stimulus B (original non-target), with positive values indicating preference for the Stimulus A.

To determine whether these comparative indices were significant for individual neurons, we used a permutation test. We shuffled the labels for each trial and recomputed the index 2000 times to create a distribution of indices that could have arisen by chance. Indices outside the center 95% interval of this distribution were considered significant (p < 0.05).

### Error trial analysis

Data from error trials were taken from behavioral sessions and imaging fields in which the mouse committed at least five false alarms. Because mice had a bias towards licking, the number of Miss trials was small, and we focused our analysis on False Alarm trials. Error modulation index (**Figure 3C**, **Figure 10C**) was computed using a renormalized auROC index as above, but by comparing stimulus period responses on False Alarm trials versus Correct Reject trials, with positive values indicating preference for False Alarm trials.

Selectivity for stimulus and choice (**Figure 3E-H**, **Figure 8C-F**) were computed using an ROC-based analysis across different time bins (Hernandez et al., 2010; Bennur and Gold, 2011). For stimulus selectivity, Hit trials were compared with False Alarm trials. For choice selectivity, False Alarm trials were compared with Correct Reject trials (this is equivalent to the error modulation index, but computed separately for each time bin). The auROC was computed by comparing the two distributions of responses at each time point (200 ms bins) during the trial, and then renormalized to range from -1 to 1. A permutation test was used to determine whether individual neurons were significantly selective (p < 0.05, 2000 permutations), and bootstrapping across imaging fields was used to determine confidence intervals on the percentage of significant neurons.

### Generalized linear model: using task predictors only

We used a generalized linear model (GLM) to regress recorded calcium signals against a time series of task events (Miri et al., 2011; Pinto and Dan, 2015; Chen et al., 2016). Calcium responses for each cell were Z-scored and modeled as the linear combination of various task events, each convolved with a filter:

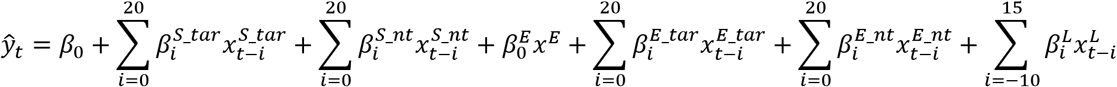

The response of a neuron at frame *t* is modelled 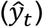 by the sum of a bias term (*β*_0_) and the weighted (*β*_i_) sum of various additional binary predictors at different lags (*i*). Binary predictors for the target stimulus 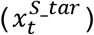 and non-target stimulus 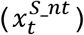 indicated the onset of stimulus presentation in either engaged or passive trials. Binary predictors for engagement included a constant offset (*x^E^*) that was 1 during engaged trials and 0 otherwise, as well as stimulus predictors 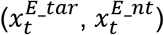 that indicated the duration of stimulus presentation during engaged trials only. Binary predictors 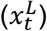 for licking indicated the duration of lick bouts, which were defined as groups of licks with an inter-lick interval less than 1 second. The number of lags were chosen to capture both the duration of the event (2 s for stimulus, 1 s for licking) as well as the slow offset dynamics of the calcium response (Chen et al., 2013b). Lags were chosen to be strictly positive (causal) for stimulus and engagement predictors, but both positive and negative (anti-causal) for licking predictors. The final model had 112 coefficients including a constant bias term. Models were fit for all task-responsive neurons, where the first 5 s from each trial (following auditory cue onset) was extracted and concatenated for analysis.

Models were fit using ridge regression using procedures similar to that of previous studies (Huth et al., 2012; Pinto and Dan, 2015; Chen et al., 2016). We first set aside 20% of trials from each condition (Hit, Correct Reject, Miss, False Alarm, Passive Target, Passive Nontarget) for testing. A regularization parameter *λ* was estimated for each cell from among a range of *λ* values (10^-2^ to 10^4^) using cross-validation. Model performance was measured by computing the proportion of explained variance, or the coefficient of determination (R^2^). Five-fold cross-validation was used to choose the *λ* that maximized the performance on the remaining 80% of training trials. The final model for each cell was fit using the best *λ,* and then performance was evaluated by measuring R^2^ of predictions for the holdout test set.

The strength of each model component (stimulus, engagement, motor/licking) was evaluated for each V1 and PPC cell (**Figure 4F**) by setting all other model coefficients to zero and finding the model predictions using just one component, averaging across both training and test trials. Model components were therefore in units of z-scored calcium response, and can be directly compared with calcium response waveforms. For stimulus and engagement components, the preferred trial type (Hit or Correct Reject) was used for each cell, whereas for the motor component, Hit trials were used for all cells. To evaluate the relative strength of the various components across V1 and PPC cells, the component strength for each cell was averaged over a window of 0 to 4 s after stimulus onset, and then a Wilcoxon rank-sum test was applied.

### Signal and noise correlations

Signal correlations between pairs of simultaneously imaged neurons were calculated as the Pearson correlation coefficient (CC) between trial-averaged responses. Noise correlations were computed by first subtracting the trial average from single trial responses and then computing the Pearson CC between the mean-subtracted responses (Hofer et al., 2011). Correlations were computed separately for Engaged and Passive conditions, and using only correct trials for the Engaged condition. Box plots (**Figure 5E**) were generated by finding, for each significantly task-responsive cell, the median Engaged noise correlation with all other simultaneously recorded cells (whether task-responsive and not) with either the same or opposite stimulus preference. The distance dependence of noise correlations (**Figure 5F**) was computed by binning all pairs of simultaneously recorded neurons (with the same stimulus preference) by their Euclidean distance in 20 μm bins up to 300 μm, and then taking the average within each bin.

### Generalized linear model: using task and network predictors

The task-only GLM can be re-written in a simplified matrix description. It models the responses of a population of *n* neurons across *t* time bins with *p* predictors as:

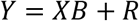

Where *Y* is a *t×n* matrix of z-scored calcium responses, *X* is a *t×p* matrix of predictors, *B* is a *p×n* matrix of task-related coefficients and *R* is a *t×n* matrix of residuals. The model prediction for the task-only GLM is simply 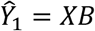.

In order to incorporate network predictors in the GLM, we considered two possible approaches. A simultaneous approach (Truccolo et al., 2005; Pillow et al., 2008) would incorporate the network terms as an additional term in the above equation:

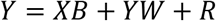

Where *W* is a *n×n* matrix of inter-neuronal network coupling weights, with the diagonal of *W* constrained to be equal to 0. In such a model, weights for task and network predictors would directly compete against each other. Given the presence of high signal correlations between neurons, this model would dramatically reduce the task weights, and also make interpretation of network weights difficult.

An alternative approach is a sequential model (Malik et al., 2011), in which network weights were estimated using the residuals of the task model:

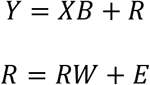

Where *W* is again a *n×n* matrix of inter-neuronal network coupling weights with a zero diagonal, and *E* is the remaining error. The final prediction can thus be computed by summing the outputs of the two models:

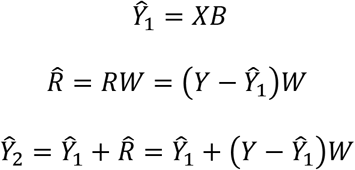

In this sequential model (**Figure 5G-J**), network weights can be naturally related to noise correlations, as both represent residual trial-to-trial fluctuations after “signal” components have been removed.

To fit the “task + network” model, the same training trials (consisting 80% of the data) were used as for the “task-only” model. Residuals for each neuron were computed by subtracting model predictions from the training data. We z-scored the residuals by dividing by the standard deviation to normalize all network weights, but kept the normalization factor for each cell in memory. For each cell, the simultaneous (lag = 0) residual activity of all other cells in the network *(R)* were used as predictors. We avoided using additional lags to keep model complexity low, which we considered reasonable given the low time-resolution of calcium responses (200 ms). Models were fit using ridge regression, as with the task-only model. We computed a single regularization coefficient *λ* for each population of cells, estimated using a random 20% subset of the cells to save computation time. Prediction performance was not significantly improved using individual values of *λ* for each cell for a handful of test populations (data not shown). The *λ* that maximized the average performance across the random subset of cells used for five-fold cross-validation was then used to fit the network model for all cells.

Final prediction performance was evaluated on the test dataset. Residuals were generated by subtracting the predictions of the task-only model from the test data 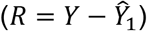. The residuals of a given cell were then predicted using the network model and the residuals of other cells in the network 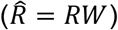. The predicted residuals for each cell were multiplied by the normalization constant and the summed with the output of the task-only model 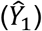 to generate final model prediction 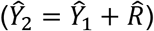.

To assess the relationship of model performance to noise correlations (**Figure 5J**), we binned neurons based on their median absolute noise correlation with other cells (bin size 0.05) and averaged the model performance within each bin.

### Contrast task analysis

For data acquired during the variable contrast task, neurons were considered significantly responsive if the mean *ΔF/F* during the stimulus period was significantly above threshold for at least two of the six contrasts of the same stimulus. We focused our analyses on target-preferring neurons, which were included if their mean response across contrasts was greater for Hit (target) trials compared to Correct Reject trials.

Single neuron contrast response functions were fit to the hyperbolic ratio function, also known as the Naka-Rushton function (Albrecht and Hamilton, 1982):

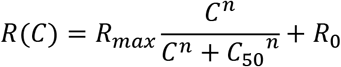

where *R(C)* is the neural response as a function of contrast, *R_max_* is the saturation point, *C_50_* is the contrast at the half-saturation point, *R_0_* is the baseline response, and *n* is an exponent that determines the steepness of the curve. The responses for both Engaged and Passive conditions were fit simultaneously, with *n* constrained to be constant across conditions, by minimizing the sum (across data points) of the squared error between the model and the data, divided by the variance of that data point.

We evaluated the goodness of fit for each neuron using a bootstrap hypothesis test, as detailed by others (Carandini et al., 1997). Briefly, we tested the null hypothesis that the mean of the probability distribution underlying the neural responses was identical to the predictions of the model. We measured the observed prediction error (*e_obs_*) and computed the probability of observing an error at least as large if the null hypothesis were true. To sample from a distribution that conformed to the null hypothesis, we shifted the data such that the mean responses equaled the model predictions, and drew 1000 bootstrap samples from this dataset, computing a prediction error (*e_i_*) for each. The proportion of samples for which the prediction error was larger than *e_obs_* is the achieved significance level. For neurons with an achieved significance level below 10% (p<0.1), there was sufficiently strong evidence against the model, and therefore these neurons were excluded from further analysis.

Contrast modulation index was computed to compare Engaged trial responses on high (64%, 32%) versus low (2%, 4%) contrasts, using a renormalized ROC index which ranged from -1 to 1, as described above. Engagement modulation index was computed by comparing Engaged versus Passive trials, using high contrast trials only. A permutation test was used to assess significance (p<0.05) of these indices by shuffling trial labels 2000 times, and comparing the measured index to the shuffled distribution of indices.

To assess anatomical spatial organization, pairwise Euclidean distance was measured between all task-responsive, target-preferring neurons in the same imaged volume (**Figure 7E-G**). Cells were grouped into four groups based on whether they were contrast-modulated, engagement-modulated only, both, or neither, and then the average within-group and across-group distance was computed for each cell (**Figure 7F**). To compute differences in modulation index as function of distance (**Figure 7G**), pairs of cells were binned by distance in 40 μm bins.

Selectivity for stimulus and choice (**Figure 8C-F**) was computed by comparing FA trials to Hit and CR trials using a frame-by-frame ROC analysis, as described above. The contrast-dependence of the auROC-based index was estimated by regressing the index against log contrast, and then finding the slope. The significance (p<0.05) of the slope was estimated for each cell by shuffling the trial labels 2000 times for each contrast and then computing the auROC on the shuffled data to generate a distribution of 2000 slopes.

### Reverse contingency task analysis

We imaged from the same field of neurons before and after reversal of reward contingency. The two sessions were separated by an average time interval of 16 ± 1 days. A semi-automated method was used to align ROIs between the two sessions (see Image Analysis). Neurons were included for analysis only if a significant response (p < 0.01) to either stimulus was observed both before and after reversal. Selectivity index was computed separately before and after reversal training (**Figure 9E-H**) by comparing responses to the Stimulus A (original target) and Stimulus B (original non-target), with positive values indicating preference for the Stimulus A. Separate permutation tests (2000 iterations) were used to assess significance of the selectivity before and after reversal. Neurons with significant selectivity (p<0.05) both before and after reversal were categorized (**Figure 10A**) based on the sign of the selectivity, and whether it was stable or altered with reversal. Engagement modulation index (**Figure 10B**) for each neuron was computed separately before and after reversal, comparing Engaged and Passive responses using the neuron’s preferred stimulus, and then taking the mean of the two values. Error modulation index (**Figure 10C**) for each neuron was also computed separately before and after reversal, comparing FA and CR responses, and then taking the mean of the two values.

## Results

### Imaging calcium responses in V1 and PPC during engaged task performance and passive viewing

We trained mice on a head-fixed lick/no-lick visual discrimination task (**Figure 1A, B**), similar to previous designs (Andermann et al., 2010; Pinto et al., 2013; Goard et al., 2016). Water-restricted mice discriminated between a target stimulus (horizontal grating drifting upwards, 0° from vertical, Stimulus A) which was rewarded with water, and a non-target stimulus of an orthogonal orientation (vertical drifting upwards rightwards, 90°, Stimulus B). Lick responses to the non-target grating were discouraged by punishment with a small, aversive drop of quinine. A retractable lick spout was presented immediately after stimulus presentation (2 s), and retracted on every trial. This restricted the animal’s lick response to the “response” period (1.5 s), and allowed us to separately assess perception and action (**Figure 1B**). Video recording of the mice during the stimulus period confirmed that mice withheld licking until the spout became available during the response epoch (data not shown). Mice (*n* = 15) achieved high levels of performance on the task (**Figure 1C**; d-prime, 2.23 ± 0.18; mean ± SEM), with a bias towards licking, resulting in more false alarms than misses.

**Figure 1.**
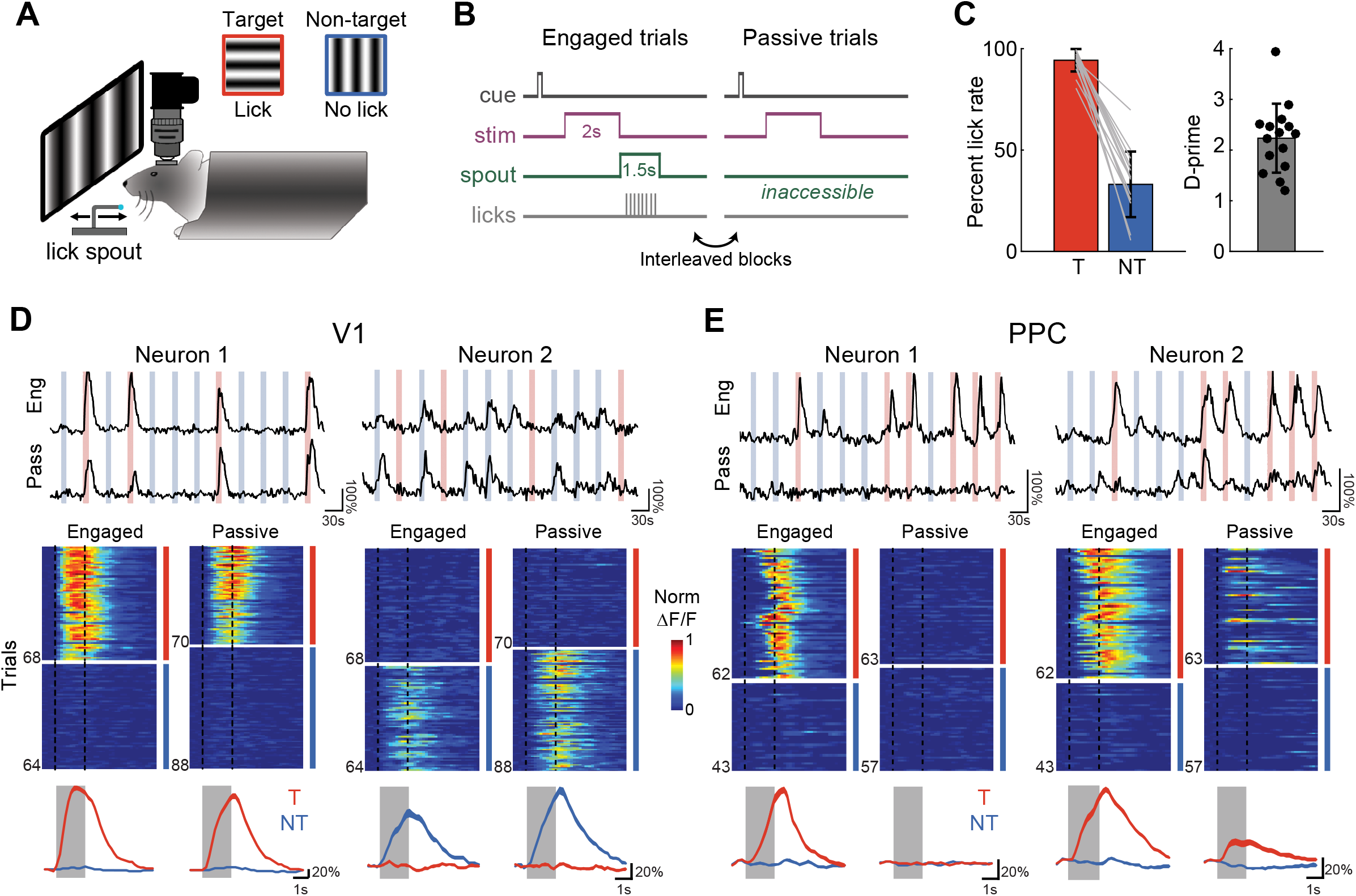
Imaging calcium responses in V1 and PPC during engaged task performance and passive viewing. (**A**) Head-fixed mice were trained to perform a go, no-go lick-based visual discrimination task. A retractable lick spout was used to restrict lick responses to a specific epoch of the task. Licks following a target stimulus (red, horizontal drifting upwards, Stimulus A) were rewarded with water, while licks to non-target stimulus (blue, vertical drifting rightwards, Stimulus B) were punished with quinine. (**B**) Trial structure for Engaged and Passive conditions. After a brief auditory preparatory cue, a drifting grating was presented for 2 s. During Engaged trials, the retractable lick spout was presented immediately after stimulus offset for a minimum of 1.5 s. During Passive trials, the spout was inaccessible. Engaged and Passive trials were presented in blocks which were usually interleaved. (**C**) Rate of licking on target (T, red) and non-target (NT, blue) for each mouse using in imaging experiments (n=15). Behavioral performance was quantified as d-prime (mean across mice: 2.22). Error bars in this and all subsequent figures depict mean ± SEM. (**D**) Stimulus-evoked response of two V1 neurons, one target-selective (left column), and one nontarget selective (right). Top, raw calcium response to multiple presentations of target (red) and non-target (blue) stimuli during both Engaged and Passive conditions. Middle, heatmap of trial-to-trial responses to target and non-target stimuli, presented in alternating blocks of Engaged (left) and Passive (right) trials, normalized to max response. Light gray shaded regions in this and subsequent figures demarcates duration of stimulus. Bottom, overlay of trial-averaged responses for each stimulus during Engaged (left) and Passive (right) conditions. Line thickness indicates mean ± SEM. (**E**) Same as (**D**) but for two PPC neurons.

We have previously demonstrated using a version of this task with a 4 s delay period that inactivation of either V1 or PPC during the stimulus period disrupts behavioral performance, whereas inactivation during later periods has no effect (Goard et al., 2016). Stimulus-period activity in PPC is therefore necessary for the task, but it is unclear whether such activity is important for sensory processing, decision formation, or action selection. In order to distinguish between these possibilities, we measured neural activity in PPC and V1 under two different conditions: during engagement in the behavioral task as well as during passive visual stimulation (**Figure 1B**). We reasoned that neurons important for sensory processing would show a robust response to stimuli during both passive and engaged conditions, whereas decision-or action-encoding neurons would show strong modulation by behavioral state.

We used two-photon microscopy and a volumetric imaging approach to image hundreds of neurons simultaneously in either V1 or PPC (see **Materials and Methods**). After behavioral training, we injected AAV2/1 syn-GCaMP6s (Chen et al., 2013b) into the two areas under stereotaxic guidance. We used a resonant scanning system combined with a z-piezo to concurrently record activity from several hundreds of GCaMP6-infected cells within layer 2/3 in a volume comprising four planes (850μm x 850μm) 20μm apart in depth, at an overall stack rate of 5 Hz. Images were corrected for X-Y movement, and fluorescence traces were extracted from semi-automatically generated ROIs based on a pixel-wise activity map (see **Materials and Methods**).

To investigate the effects of behavioral performance on neural responses, we imaged the same neurons in alternating blocks (5-10 min, 40-80 trials) of engaged behavior and passive viewing (**Figure 1B**). During “engaged” behavior trials, the spout was extended on each trial during the response period. During “passive” trials, the spout was not extended during the response period but remained withdrawn and inaccessible from the animal. To avoid extraneous stimulus confounds, no additional cue was provided to signal engaged or passive blocks. Nonetheless, mice rapidly became aware after the first 1-2 trials of a block whether the spout would be available for a behavioral response, as confirmed in video recordings by the complete lack of licking during passive blocks (data not shown). Discrimination performance was similarly high for both the first and subsequent “engaged” blocks.

We imaged from an average of 606 cells (range: 257-1057) in each population within either V1 or PPC, of which an average 135 cells (range: 21-364) exhibited significant task-related responses. Pilot experiments using transgenic mice with tdTomato expressed in PV+ and SOM+ interneurons revealed that calcium signals from these neurons were generally too weak to be measured with volume scanning, and therefore the vast majority of task-responsive cells were likely to be excitatory pyramidal neurons (data not shown). We first analyzed only the neural responses measured during correct Engaged trials, or during Passive viewing. In V1, many neurons showed a robust and reliable response to either the target or non-target stimulus, though some were modulated by engagement in the task. For example, one target-preferring cell showed a reliable passive response to the target stimulus that was moderately enhanced during task performance (**Figure 1D**, left). A non-target preferring cell, however, exhibited a suppressed response during engagement in the task (**Figure 1D**, right). By contrast, PPC neurons exhibited much stronger responses during task engagement compared to passive viewing, specifically for target stimuli. For example, one target-preferring cell had robust activity only during engagement (**Figure 1E**, left), whereas another cell showed relatively weak passive responses that became stronger during task performance (**Figure 1E**, right).

### V1 neurons respond passively, whereas PPC responses are gated by task engagement

To compare the overall response properties of V1 and PPC, we analyzed the responses of all neurons that had significant stimulus-period activity during the task. We focused on stimulus-period responses given that inactivation during this period disrupts behavioral performance (Goard et al., 2016). A total of 1915 neurons (18% of all neurons, 18 fields, 9 mice) in V1 (**Figure 2A, B**) and 3524 neurons (26% of all neurons, 22 fields, 10 mice) in PPC (**Figure 2C, D**) were significantly responsive during the task. Two striking differences between V1 and PPC were immediately apparent when examining the trial-averaged responses. First, while V1 was about evenly split between target- and nontarget-preferring cells (60.6% ± 5.8% target-preferring, **Figure 2E**), there was a stronger bias in PPC towards the target stimulus (87.6% ± 7.4% target-preferring; V1 versus PPC, p = 4.68 × 10^-5^, Wilcoxon rank-sum test). Secondly, while most task-responsive V1 neurons also responded during passive viewing (88.3% ± 6.8% with significant passive response, **Figure 2F**), the majority of PPC neurons had responses that were gated by task engagement (only 29.7% ± 10.3% with significant passive response; V1 versus PPC, p = 4.71 × 10^-7^, Wilcoxon rank-sum test).

**Figure 2.**
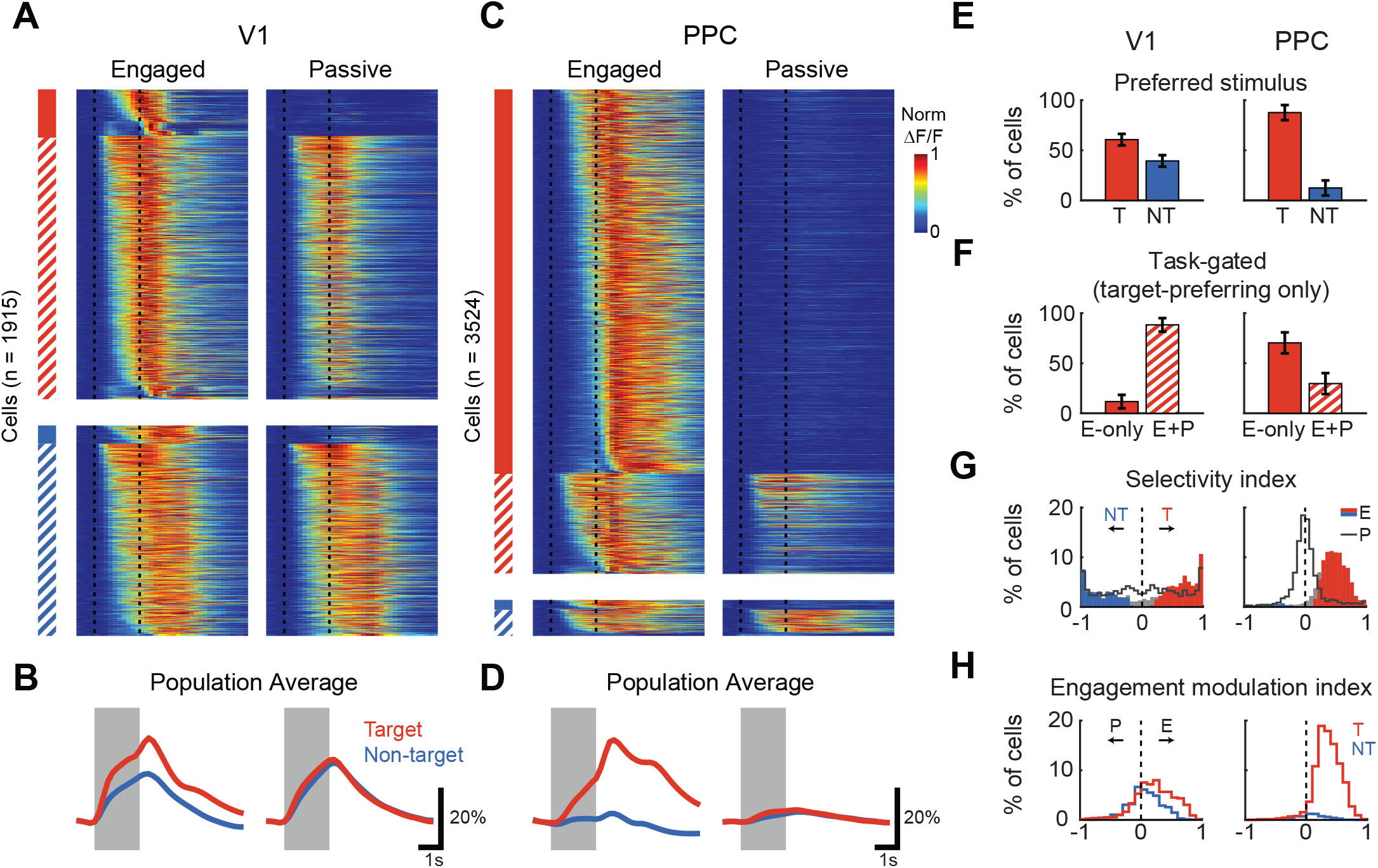
V1 neurons respond passively, whereas PPC responses are gated by task engagement. (**A**) Trial-averaged responses of all task-responsive V1 neurons (n = 1915). Heatmap of all trial-averaged responses (preferred stimulus only) in both Engaged (left) and Passive (right) conditions, normalized by peak response. Neurons are separated into target-preferring (red) and non-target preferring (blue), and then into passive-responding (solid) and task-gated (hatched). Vertical dashed lines demarcate duration of stimulus. (**B**) Bottom, average response across V1 neurons to the target (red) and non-target (blue) in Engaged (left) and Passive (right) conditions. Line thickness indicates mean ± SEM. (C-D) Same as (A-B), but for PPC neurons (n = 3524). (**E**) Percentage of neurons in V1 (left) and PPC (right) that prefer target (T) or non-target (NT) stimulus. Error bars represent bootstrapped SEM across sessions. (**F**) Percentage of target-preferring neurons in V1 (left) and PPC (right) that are task-gated, i.e. that respond only during engagement (E-only), or that respond during both engaged and passive conditions (E+P). (**G**) Histogram of stimulus selectivity index for V1 (left) and PPC (right) during Engaged (filled bars) and Passive conditions (gray line). Positive selectivity indicates preference for Target. Colored bars indicate neurons with significant individual selectivity during Engaged trials. (**H**) Histogram of engagement modulation index for V1 (left) and PPC (right) for target-(red) and nontarget-preferring (blue) neurons.

The dramatic effect of task engagement on PPC responses can also be observed by comparing the selectivity of responses during Passive and Engaged conditions (**Figure 2G**). We quantified selectivity using an ROC-based index that ranged from -1 to 1, with positive values indicating preference for the target stimulus (see **Materials and Methods** for details). V1 has a large proportion of significantly selective (p < 0.05, permutation test) neurons in Passive conditions (85.3% ± 3.8% of cells), with a small but significant increase in Engaged conditions (91.8% ± 2.5% of cells; Passive vs Engaged, p = 4.97× 10^-3^, Wilcoxon signed-rank test). By contrast, PPC neurons are largely unselective during Passive conditions (41.9% ± 10.8% selective), and yet become significantly selective to target trials during task engagement (89.5% ± 4.7% of cells; Passive vs Engaged, p = 5.95 × 10^-5^, Wilcoxon signed-rank test). We also quantified the change in responses for engagement versus passive viewing for each neuron using an engagement modulation index (**Figure 2H**), which also ranged from -1 to 1, with positive values indicating increases with engagement. The mean modulation index was significantly above zero for target-preferring neurons in both V1 (0.227 ± 0.009, n = 1148; p < 5.0 × 10^-4^, permutation test, 2000 permutations) and PPC (0.351 ± 0.004, n = 3292; p < 5.0 × 10^-4^). Nontarget-preferring neurons in V1 showed weaker modulation (modulation index: 0.059 ± 0.010 for non-target, n = 767) than target-preferring neurons (p = 9.07 × 10^-4^, clustered Wilcoxon rank-sum test), indicating that the effect of engagement was stimulus-specific.

Comparing responses during engaged behavior and passive viewing therefore revealed different response properties in V1 and PPC. In V1, task performance did not merely increase overall responsiveness (as would be expected with arousal), but instead modulated firing rates to enhance the contrast between target and non-target stimuli. By contrast, PPC responses were strongly target-selective and gated by behavior. This dramatic task-dependent increase in activity strongly implicates a role for PPC beyond mere sensory processing. Furthermore, because this activity is selective to target trials in which the animal licks, a major component of PPC responses likely signals the impending action and does not simply reflect overall increases in arousal or attention during task engagement. A subset (~30%) of PPC neurons, however, do have significant passive responses, and could play a role in sensory processing.

### Error trials reveal sensitivity of PPC to both stimulus and choice

One possible explanation for the strong behavioral gating and target selectivity of PPC responses is that these neurons reflect signals related to movement or action planning. Another possibility is that task engagement provides a stimulus-specific attentional signal. One way to arbitrate between these alternatives is to observe activity level on error trials. Movement-related activity would appear on error trials when the animal moves inappropriately, whereas stimulus-specific signals would only appear on target stimulus trials. We thus examined PPC responses during error trials to see whether activity in PPC reflected the stimulus, the animal’s eventual choice, or both. We focused on target-selective neurons, as these represented the vast majority of responses in PPC, and restricted analysis to behavioral sessions and imaging fields in which the mouse committed at least five False Alarm (FA) trials (V1, n = 1053 cells from 16 fields; PPC, n = 3034 cells from 21 fields). Although similar analyses could be performed using Miss trials, mice were biased towards licking and usually had very low Miss rates.

For V1, individual target-preferring neurons varied in their responsiveness on nontarget trials (**Figure 3A**, left), with a subset of neurons showing significant nontarget responses. However, the average strength of the nontarget response was similar on FA trials compared to Correct Reject (CR) trials, regardless of the animal’s decision to lick (**Figure 3B**, left). By contrast, PPC neurons invariably showed stronger responses on FA versus CR trials (**Figure 3A**, right), as also seen in the population average (**Figure 3B**, right). We quantified the modulation of each neuron’s response (measured during the stimulus period) on FA versus CR trials using an error modulation index that ranged from -1 to 1, with positive values indicating stronger FA responses (**Figure 3C**). The mean error modulation index was significantly above zero for PPC (0.109 ± 0.003, p = 0.014, clustered Wilcoxon signed-rank test) but not for V1 (0.014 ± 0.006, p = 0.462). However, the overall difference between PPC and V1 did not reach significance (p = 0.169, clustered Wilcoxon rank-sum test), and the fraction of significantly error-modulated neurons was not significantly larger in PPC (13.8% ± 7.3%) than in V1 (7.5% ± 4.2%; p = 0.149, Wilcoxon rank-sum test). The mean stimulus-period response of PPC neurons is therefore only weakly modulated by the animal’s impending choice.

**Figure 3.**
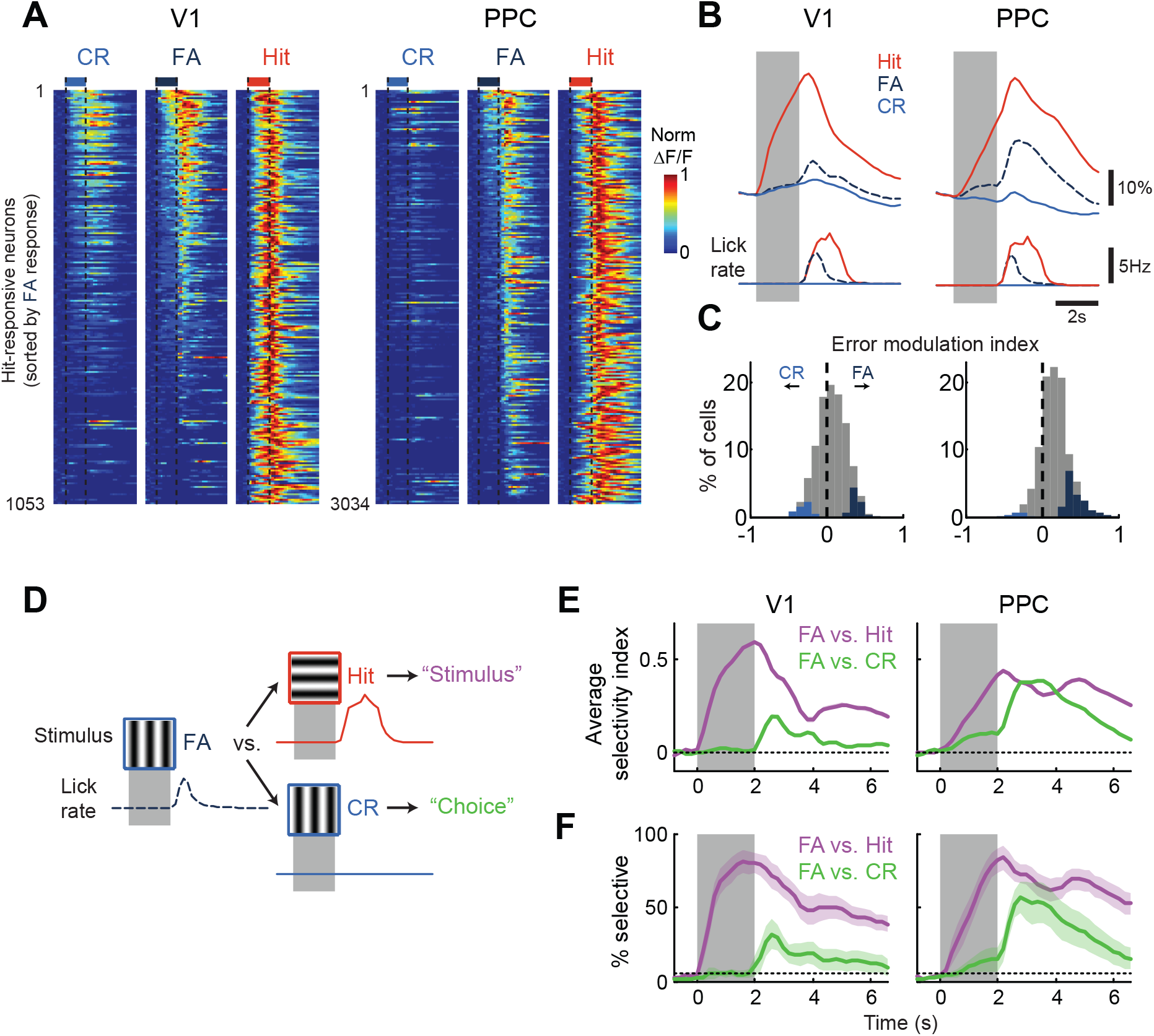
Error trials reveal sensitivity of PPC to both stimulus and choice. (**A**) Trial-averaged responses during False Alarm (FA) trials, for recordings with at least five FA trials, in V1 (left; 1053 neurons across 16 sessions) and in PPC (right; 3034 neurons across 21 sessions). For each heatmap, responses of all Hit-preferring neurons during Correct Reject (CR, left, blue), FA (center, dark blue), and Hit trials (right, red) are normalized by peak response. Neurons are sorted by descending strength of FA response. Vertical dashed lines demarcate duration of stimulus. (**B**) Population trial-averaged responses (top) during Hit, FA (dashed), and CR trials, for V1 (left) and PPC (right). Average lick rate (bottom). (**C**) Histogram of error modulation index for V1 (left) and PPC (right) neurons, computed by comparing responses on FA and CR trials during the stimulus period using an ROC-based analysis. Colored bars indicate neurons with significant individual modulation (p < 0.05). (**D**) Using a time-dependent ROC-based analysis, FA trials are compared with Hit trials to assess “Stimulus” selectivity (purple), and are compared with CR trials to assess “Choice” selectivity (green). (**E**) Average selectivity measured using a frame-by-frame ROC analysis. Selectivity can range from -1 to 1, with positive values indicating stronger Hit responses (green) or weaker CR responses (purple), as compared to FA responses. Line thickness indicates mean ± SEM. (**F**) Percentage of neurons with significant (p < 0.05) positive selectivity. Shading indicates mean ± SEM across sessions. Dotted line indicates the expected percentage by chance.

Averaging across the stimulus period, however, could obscure differences in the dynamics of stimulus and choice information in the two areas. We therefore also performed a time-dependent error analysis (Goard et al., 2016). We used an ROC-based analysis to compare responses on FA trials with responses on correct trials at each time point of the trial (Britten et al., 1996; Hernandez et al., 2010). Using only target-preferring neurons, we could assess selectivity for the stimulus by comparing FA to Hit trials, and selectivity for the choice by comparing FA to CR trials (**Figure 3D**). Selectivity for each neuron ranged from -1 to 1, with positive values indicating stronger Hit responses or weaker CR responses, as compared to FA trials. We quantified both the average selectivity across neurons (**Figure 3E**), and the proportion of neurons with significant selectivity (**Figure 3F**).

In V1, stimulus selectivity was strong and appeared rapidly after stimulus presentation (**Figure 3E**, left; significantly different from 0, p < 0.05 from t = 0.2 s to 6.6 s, clustered Wilcoxon signed-rank test), and then gradually declined due to the slow decay of the GCaMP6s indicator. A majority of V1 neurons showed significant selectivity between FA and Hit trials within the stimulus period (84.0% ± 6.6%; **Figure 3F**, left). Choice selectivity was weak, with average selectivity close to zero during the stimulus period, and only becoming significant after stimulus offset (p < 0.05 from t = 2.4 s to 4.0 s, clustered Wilcoxon signed-rank test). Similarly, the proportion of significantly-selective neurons (bootstrapped across imaging fields) was no greater than the chance level of 5% until after stimulus offset. For PPC, however, we observed significant selectivity for both stimulus (p < 0.05 from t = 0.2 s to 6.6 s) and choice (p < 0.05 from t = 0.8 s to 6.0 s) within the stimulus period (**Figure 3E**, right). Most PPC neurons had significant stimulus selectivity (73.4% ± 9.7%), showing stronger responses on Hit versus FA trials. Choice selectivity peaked later in the trial, but the proportion of significantly-selective neurons was above chance as early as 1.0 s after stimulus onset. These results suggest that PPC activity is neither purely sensory, as V1 responses appear to be, nor is it purely movement-related. Instead, PPC appears to be sensitive to both the stimulus and impending choice of the animal.

### Generalized linear model (GLM) of V1 and PPC responses

PPC responses are sensitive to stimulus, choice, and engagement, but can these various components be dissociated? We used a generalized linear model (GLM) (Miri et al., 2011; Pinto and Dan, 2015; Chen et al., 2016) to disambiguate the contributions of stimulus, engagement, and motor action on single neuron calcium responses across both Passive and Engaged conditions. Responses of each V1 and PPC cell were modeled (see **Materials and Methods**) as a linear combination of components that were time-locked to stimulus presentation or licking onset (**Figure 4**). We quantified the performance of the GLM as the proportion of variance explained (R^2^) for a separate test dataset not used for fitting (**Figure 4E**). Using all significantly task-responsive neurons, the average model performance was higher for V1 (0.334 ± 0.005, n = 1915 cells) than for PPC (0.190 ± 0.003, n = 3524; V1 versus PPC, p = 4.01 × 10^-5^, clustered Wilcoxon rank-sum test).

**Figure 4.**
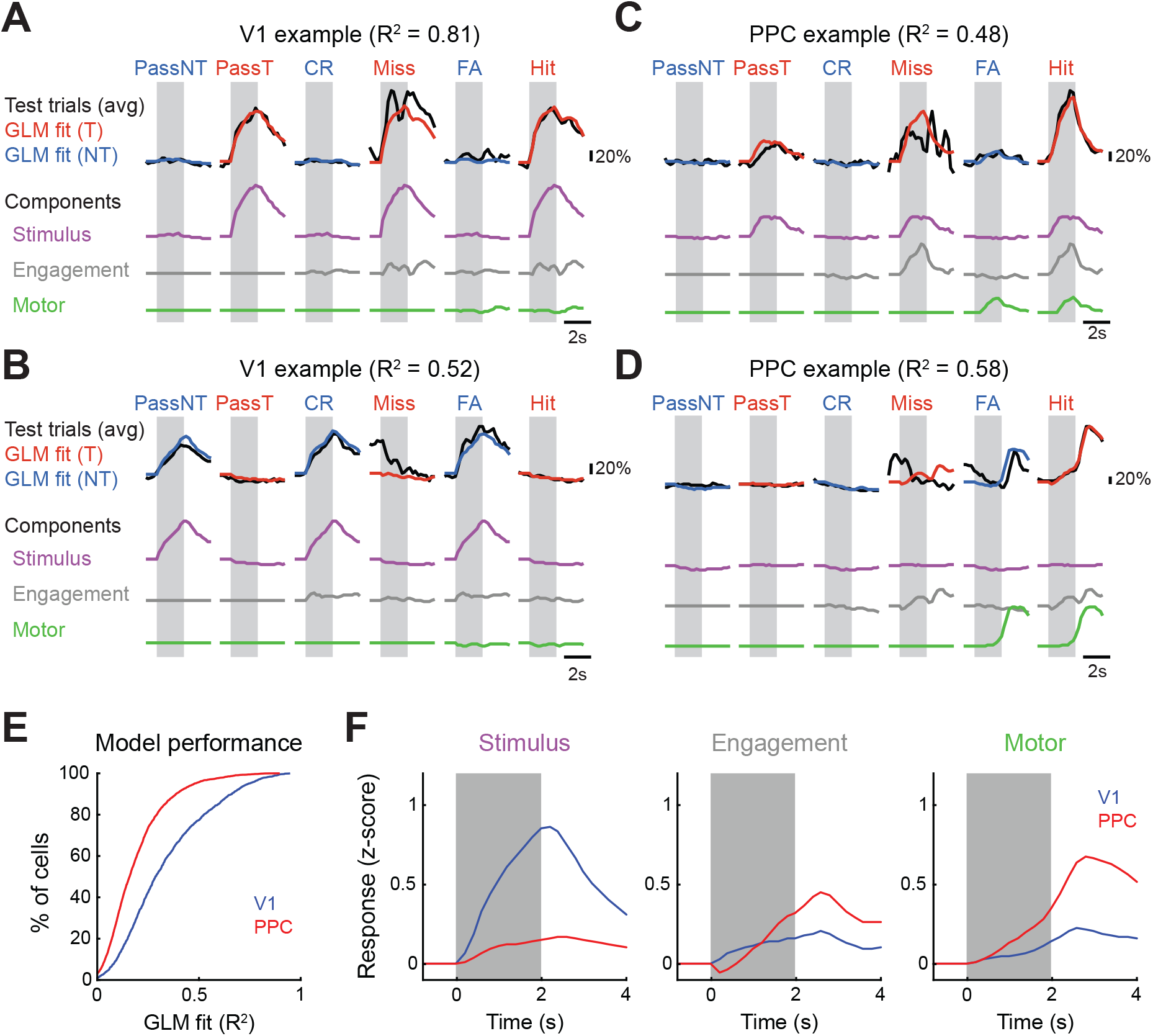
Generalized linear model (GLM) of V1 and PPC responses. A generalized linear model was fit to training data to predict the calcium responses of individual V1 and PPC neurons using lagged predictors for stimulus, engagement and motor action (licking). (**A**) GLM model performance for an example V1 neuron. GLM fits on Target (red) and Non-target (blue) trials were compared with trial-averaged test data (black) across Passive (PassNT, PassT), Engaged No-Lick (Miss, FA), and Engaged Lick (FA, Hit) conditions. GLM components related to Stimulus (purple), Engagement (gray), and Motor (green) were summed to generate overall GLM fit. Shaded region represents time of stimulus presentation. (**B**) Same as (**A**), but for a different, nontarget-preferring V1 neuron. (**C-D**) GLM model performance for two example PPC neurons. One neuron has a strong Engagement component (**C**), while the other has a stronger Motor component (**D**). (**E**) Cumulative histogram of model prediction performance on test trials, quantified for each cell as fraction of variance explained (R^2^), for both V1 (blue, n = 1915 cells) and PPC (red, n = 3524). (**F**) Population-averaged GLM model components related to Stimulus (left), Engagement (middle), and Motor (right), for both V1 (blue) and PPC (red). For Stimulus and Engagement components, the preferred stimulus component for each cell was used for averaging. Calcium responses were z-scored before model fitting. Light shaded gray region demarcates duration of stimulus.

We then assessed the relative contribution of stimulus, engagement, and motor model components for cells in each area, and illustrate the performance of the model on a few examples (**Figure 4A-D**). Most V1 cells exhibited a strong stimulus component (0.494 ± 0.013, component of z-scored calcium response, see **Materials and Methods**), which reflected sensory drive on Passive trials, whether the neuron preferred the target (**Figure 4A**) or the nontarget (**Figure 4B**) stimulus. By contrast, PPC neurons had much weaker stimulus components (**Figure 4F**, left; 0.117 ± 0.013; V1 versus PPC, p = 6.84 × 10^-5^, clustered Wilcoxon rank-sum test). The engagement component of the GLM reflected stimulus-specific signals that occurred exclusively on Engaged trials. This could be disambiguated from the stimulus-independent motor component, which would be present on both Hit and False Alarm (FA) trials. Individual PPC neurons could vary in the relative contribution of engagement and motor components (**Figure 4C-D**), but on average PPC cells exhibited a slightly (but not significantly) stronger engagement component (**Figure 4F**, middle; V1, 0.183 ± 0.009; PPC, 0.233 ± 0.006; V1 versus PPC, p = 0.144, clustered Wilcoxon rank-sum test) and a much stronger motor component (**Figure 4F**, right; V1, 0.155 ± 0.005; PPC, 0.371 ± 0.005; V1 versus PPC, p = 0.015, clustered Wilcoxon rank-sum test) as compared to V1 cells. These results suggest that PPC responses reflect both stimulus-specific engagement signals and choice-related motor signals, which can be partially explained using a linear model.

### Addition of noise correlations in GLM improves model prediction, especially for PPC

The GLM was sufficient to predict response properties of neurons from both areas, but a large proportion of variance remained unexplained, particularly for PPC neurons (~20% variance explained). One possible source of unexplained variance is the fluctuation of network activity from trial-to-trial. These inter-neuronal dependencies, often termed noise correlations, can be relatively strong in sensory cortex, especially during inattentive states (Zohary et al., 1994; Cohen and Maunsell, 2009; Cohen and Kohn, 2011). We therefore quantified the strength of these noise correlations in V1 and PPC during both task performance and passive viewing.

We divided neurons from each population based on their stimulus preference during Engaged trials. Neurons with the same stimulus preference naturally exhibited large signal correlations (computed as the correlation between trial-averaged responses during correct Engaged trials) in both V1 (**Figure 5A**) and PPC (**Figure 5B**), though there was a much larger proportion of Target-preferring neurons in PPC. We then computed noise correlations in both Engaged and Passive conditions by correlating trial-to-trial fluctuations across simultaneously recorded neurons. Noise correlations between neurons with the same stimulus preference were stronger than noise correlations between pairs with opposite stimulus preferences for both V1 (**Figure 5E**; opposite, 0.040 ± 0.001; same, 0.110 ± 0.002; p = 1.94 × 10^-4^, clustered Wilcoxon signed-rank test) and PPC (opposite, 0.051 ± 0.002; same, 0.197 ± 0.002; p = 5.39 × 10^-5^). Focusing on pairs with the same stimulus preference, V1 neurons exhibited similar strength of noise correlations in Engaged versus Passive conditions (**Figure 5C**; passive, 0.109 ± 0.002; engaged, 0.110 ± 0.002; p = 0.388, clustered Wilcoxon signed-rank test). In contrast, PPC cells had stronger noise correlations in Passive versus Engaged conditions (**Figure 5D**; passive, 0.254 ± 0.002; engaged: 0.195 ± 0.002; p = 4.05 × 10^-3^). Comparing across all cells and all populations, noise correlations were stronger in PPC compared to V1 (p = 6.40 × 10^-4^, clustered Wilcoxon signed-rank test), and with the strongest correlations between pairs within 50 μm of each other (**Figure 5F**) (Rikhye and Sur, 2015). These results indicate that network correlations strongly affect neural responses, particularly in PPC.

**Figure 5.**
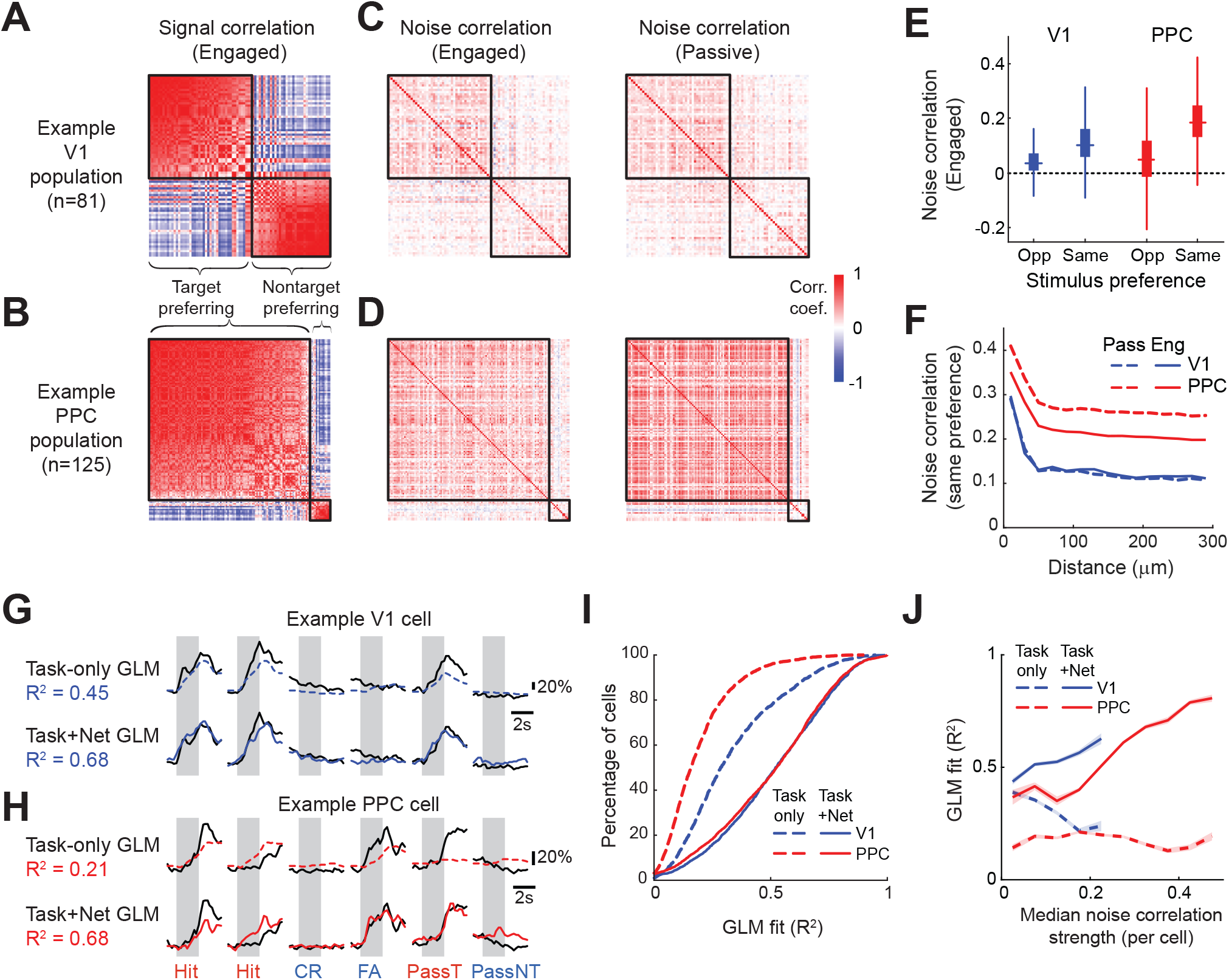
Addition of noise correlations in GLM improves model prediction, especially for PPC. (**A**) Heatmaps of signal correlations during Engaged trials in a population of V1 cells (n=81). Cells are clustered by stimulus preference, and then sorted by mean within-cluster correlation. (**B**) Same as (**A**) but for a population of PPC cells (n=125). (**C**) Heatmaps of noise correlations during Engaged (left column) and Passive (right column) trials in a population of V1 cells. Neurons are sorted in the same order as in (**A**). (**D**) Same as (**C**) but for the population of PPC cells in (**B**). (**E**) Box plot of noise correlation (during Engaged trials) in V1 (left, blue) and PPC (right, red) for pairs with the same or opposite (Opp) stimulus preference. (**F**) Average noise correlation between pairs with same stimulus preference as a function of between-cell distance in V1 and PPC during Engaged and Passive trials. (**G**) Performance of two GLM models, one using only task-related predictors (Task-only, dashed colored lines), and one using both task-related and network predictors (Task+Net, solid colored lines) for one example V1 cell. The performance of each model was assessed on the same set of test trials (six example trials shown) as the fraction of variance explained (R^2^). Shaded regions indicate stimulus presentation, and trial type is indicated with a label (Hit, CR, etc). (**H**) Same as (**G**) but for an example PPC cell. (**I**) Cumulative histogram of model prediction performance on test data, quantified for each cell as fraction of variance explained (R^2^), for both V1 (blue) and PPC (red) and for both the Task-only model (dashed, same as Figure 4E) and the Task + Network model (solid). (**J**) Task + Network model performance is correlated with the strength of noise correlations for a given cell. Cells were binned based on their median absolute noise correlations with other cells, and shading indicates SEM.

We then sought to incorporate correlations between neurons in our GLM as network coupling terms to see if they would improve prediction performance (Truccolo et al., 2005; Pillow et al., 2008). We reasoned that because of large signal correlations between neurons, directly incorporating the network terms into the original model would mask the contribution of the various task events. Therefore, we first fit the data to the original model which captured the “signal” components (Malik et al., 2011), and then fit the residual errors for each neuron using a network model (see **Materials and Methods**). We then compared the performance of the original “task-only” GLM to the “task + network” model on test data that was not used to train either model.

Although the “task-only” GLM was proficient at predicting trial-averaged responses, it failed to capture variations in responses between trials with the same behavioral outcome, e.g. two different Hit trials. However, as illustrated by example cells from V1 (**Figure 5G**) and PPC (**Figure 5H**), incorporation of network coupling terms improved the model’s ability to predict responses on single trials. Overall model prediction performance (**Figure 5I**) improved for cells in V1 (R^2^, task-only: 0.334 ± 0.005; task+net: 0.507 ± 0.006), but even more so for neurons in PPC (task-only: 0.190 ± 0.003; task+net: 0.490 ± 0.005; improvement for PPC greater than V1, p = 8.83 × 10^-5^, clustered Wilcoxon rank-sum test), as predicted by the high level of noise correlations measured in PPC responses. We further examined the relationship between model performance and the strength of noise correlations on a cell-by-cell basis (**Figure 5J**). There was a strong positive correlation for both V1 (Pearson’s correlation, r = 0.147, p = 1.07 × 10^-10^) and PPC (r = 0.382, p = 7.83 × 10^-123^), indicating that cells that were more strongly influenced by network inputs could be better explained using the “task + network” model. Interestingly, noise correlation strength was inversely correlated with task-only model performance for V1 (r = -0.228, p = 6.52 × 10^-24^), with a weaker relationship for PPC (r = -0.050, p = 3.28 × 10^-3^). This suggests that for V1 neurons, the ability of a linear model to predict responses based on task parameters (i.e. stimulus) is corrupted by the presence of noise correlations (Zohary et al., 1994). Overall these results demonstrate that both V1 and PPC responses can be better described when incorporating network correlations.

### PPC reflects both stimulus contrast and behavioral state

Previous work has shown that in both primates (Shadlen and Newsome, 2001; Gold and Shadlen, 2007) and rats (Hanks et al., 2015), neurons in PPC encode not only the impending motor action but also the sensory evidence for that decision. To test whether neurons in mouse PPC similarly reflected the decision process, we varied the amount of sensory evidence from trial-to-trial by manipulating stimulus contrast. We also compared responses during engaged and passive conditions to examine how sensory and motor signals may be combined in PPC responses.

A subset of the mice (n = 6) were trained to perform a variant of the discrimination task, in which the contrast of the grating stimulus varied randomly from trial-to-trial (**Figure 6A**). Mice performed well above chance, even at very low contrasts (d’ at 2% contrast, 1.05 ± 0.25; p = 0.031, Wilcoxon signed-rank test), although performance degraded as contrast was lowered (p = 1.10 × 10^-4^, Friedman test) reflecting the decrease in the strength of sensory evidence (**Figure 6B**). We imaged from neurons in V1 (n = 8 fields, 3 mice) and PPC (n = 11 fields, 6 mice) and computed contrast response functions for both passive and engaged conditions. We again focused our analysis on target-preferring neurons (V1, n = 250; PPC, n = 611), which constituted the vast majority of task-responsive PPC neurons (95% in this dataset). Neurons were included for further analysis if they demonstrated significant Hit responses at multiple contrasts that could be well fit with a hyperbolic ratio function (Albrecht and Hamilton, 1982) (see **Materials and Methods**).

**Figure 6.**
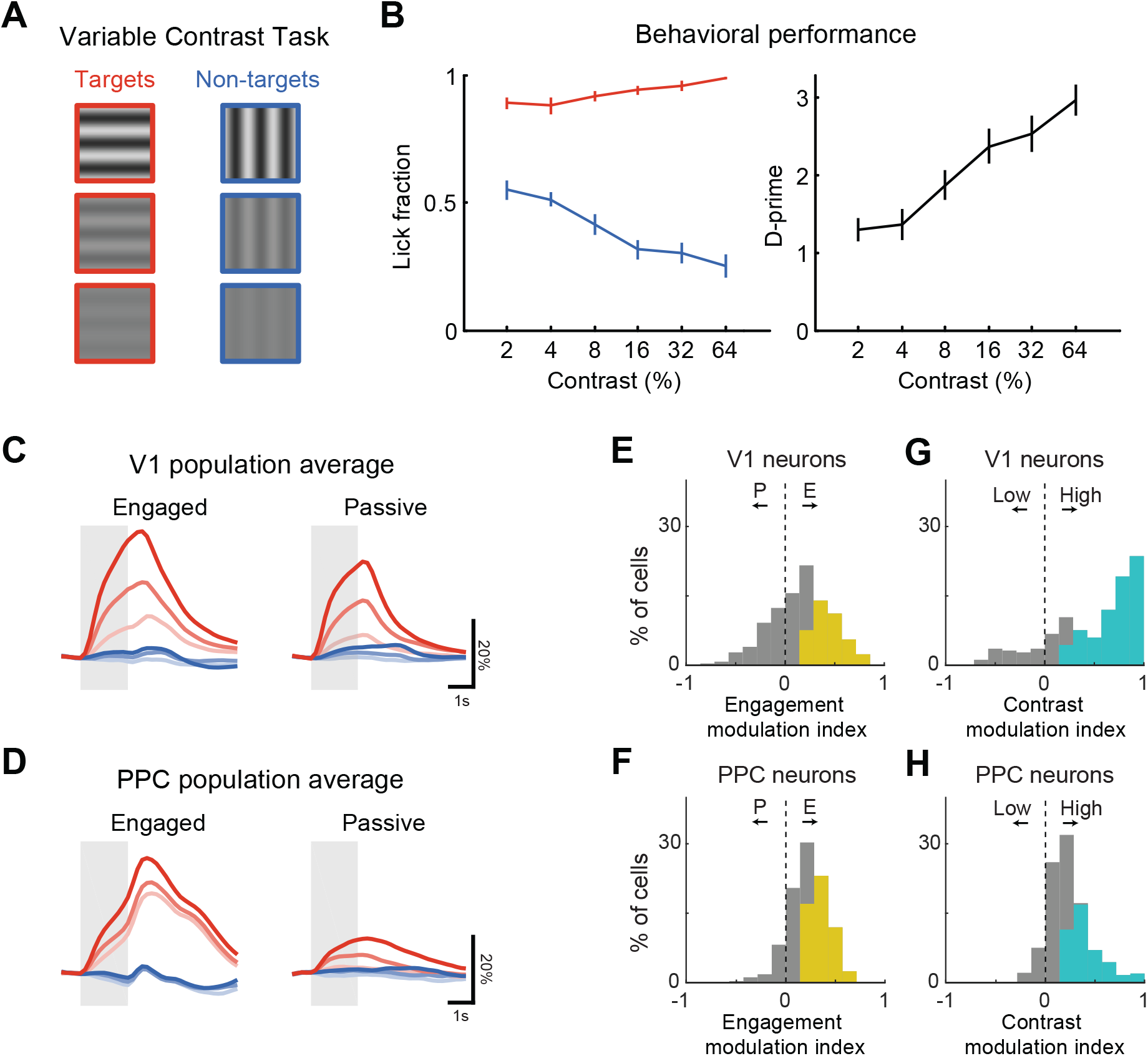
PPC reflects both stimulus contrast and behavioral state. (**A**) Mice discriminated between orthogonally oriented Target and Non-target stimuli with contrast varying from 2 to 64%. (**B**) Behavioral performance (d-prime) on the variable contrast discrimination task, averaged across 19 sessions from 6 mice. Error bars indicate SEM. (**C**) Population trial-averaged response of all target-preferring V1 neurons (n = 250) across contrasts during correct Engaged (left) and Passive (right) target (red) and nontarget (blue) trials. Responses are averaged across low (2 or 4%, light shade), medium (8 or 16%, medium shade), and high (32 or 64%, dark shade) contrast. Light gray shaded regions demarcate duration of stimulus. (**D**) Same as (**C**) but for PPC (n = 611 target-preferring neurons). (**E-F**) Histograms comparing engagement modulation in V1 (**E**) and PPC (**F**), computed by comparing responses on Engaged versus Passive high contrast trials. Colored bars indicate neurons with significant modulation. (**G-H**) Histograms comparing contrast modulation in V1 (**G**) and PPC (**H**), computed by comparing responses on high versus low contrast Engaged trials. Colored bars indicate neurons with significant modulation.

As in the single-contrast task, the population response in V1 during the variable contrast task was robust in both Engaged and Passive conditions (**Figure 6C**), with most neurons showing a significant response to at least one contrast during both conditions (68.9% ± 11.8% of cells). Conversely, PPC population activity was more robust in Engaged versus Passive conditions (**Figure 6D**), with only a subset of PPC neurons showing significant passive responses (27.8% ± 14.9%; V1 versus PPC, p = 7.59 × 10^-3^, Wilcoxon rank-sum test). We quantified the strength of modulation by contrast and engagement for each neuron using ROC-based indices that ranged from -1 to 1 (**Figure 6E-H**). Engagement modulation index was similarly high in V1 and PPC (**Figure 6E-F**; V1, 0.149 ± 0.019; PPC, 0.222 ± 0.008; V1 versus PPC, p = 0.209, clustered Wilcoxon rank-sum test), but modulation by contrast was stronger on average in V1 compared to PPC (**Figure 6G-H**; V1, 0.487 ± 0.027; PPC, 0.233 ± 0.009; V1 versus PPC, p = 0.021, clustered Wilcoxon rank-sum test). Nonetheless, the mean contrast modulation index in PPC was significantly greater than zero (p = 4.86 × 10^-3^, clustered Wilcoxon signed-rank test).

PPC is on average modulated by both engagement and stimulus contrast, but closer examination of the individual PPC contrast response functions revealed a great deal of heterogeneity. We therefore divided neurons into groups based on whether they exhibited significant modulation by contrast and/or engagement (**Figure 7**). A subset of PPC neurons (21.2% ± 13.5% of cells) showed strong modulation by contrast, but very weak modulation by engagement (**Figure 7C, D**, left column). Such neurons therefore faithfully represented the sensory stimulus regardless of behavioral state. Conversely, a larger group of PPC neurons (28.1% ± 12.3%) were gated by task engagement but showed little to no modulation with contrast (**Figure 7C, D**, middle column). These neurons reflected the behavioral state and impending action of the animal irrespective of sensory drive. Lastly, a third subset of PPC neurons (28.2% ± 10.6%) were significantly modulated by both contrast and engagement (**Figure 7C, D**, right column). PPC therefore appears to contain both contrast-modulated “sensory” neurons as well as engagement-modulated “motor” neurons. This differs from V1 (**Figure 7A**, left; p = 7.10 × 10^-17^, chi-squared test of independence), where most neurons (74.8% ± 7.0%) are modulated by contrast, and much fewer by engagement alone (9.2% ± 7.1%).

**Figure 7.**
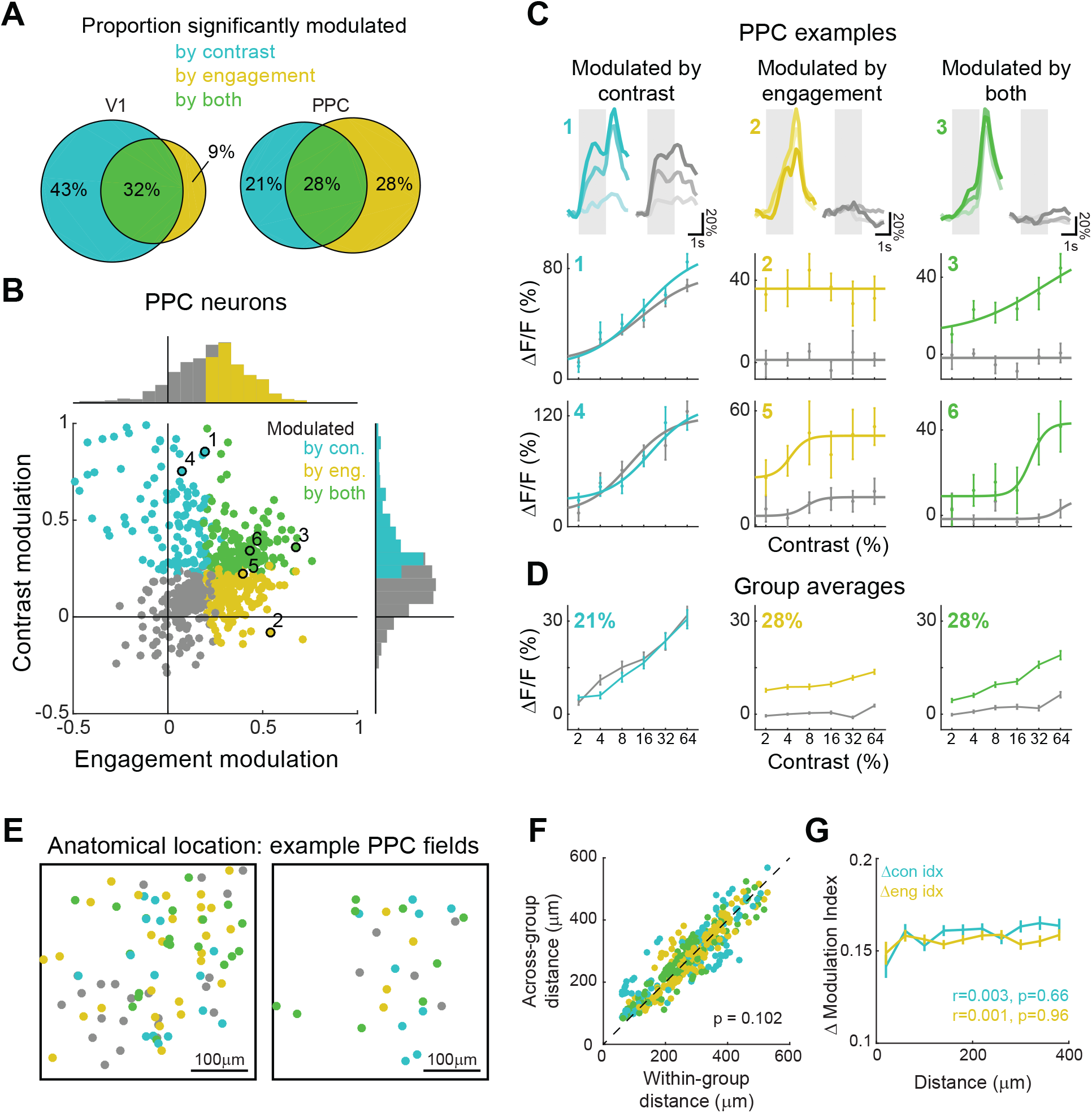
PPC neurons are heterogeneous in their sensitivity to contrast and engagement. (**A**) Venn diagrams indicating proportions of target-preferring neurons in V1 and PPC with significantly positive modulation by contrast alone (cyan), engagement alone (yellow), or both contrast and engagement (green). (**B**) Scatter plot and histograms of contrast modulation versus engagement modulation for all target-preferring PPC neurons (n = 611) imaged during the variable contrast task. Colored bars in histograms (same as Figure 5F, H) indicate neurons with significantly positive modulation by contrast (cyan) or engagement (yellow). Colored dots on scatter plot demarcate neurons with significantly positive modulation by contrast alone (cyan), engagement alone (yellow), or both contrast and engagement (green). Individual examples in (**C**) are marked with the corresponding number. (**C**) Trial-averaged responses (top row) and contrast-response functions (middle and bottom rows) of example PPC neurons that were significantly modulated by contrast (left column), engagement (middle column), or both (right column). Modulation index values for each example can be found by referring to (**B**). (**D**) Group-averaged contrast-response functions. Percentages indicates proportion of PPC neurons within each group. (**E**) Relative spatial location of neurons with modulation by contrast, engagement, or both in two example imaging sessions. Spatial locations of cells from 4 planes are collapsed into one. Scale bar, 100 μm. (**F**) Average within-group and across-group distances for each cell, colored by group. No significant difference was found for any individual group or for the whole population (p = 0.102). (**G**) Difference in engagement (yellow) and contrast (cyan) modulation indices as a function of distance between cells. Distances beyond 400 μm are not shown. No significant correlation between modulation index and distance was found.

We then considered whether there was any anatomical organization of these functional properties within PPC. For each imaged PPC volume, functional subpopulations of contrast-modulated and engagement-modulated cells appeared to be intermingled across space (**Figure 7E**). We computed the pairwise distance between all target-preferring neurons, and found no significant difference between within-group and across-group distances (**Figure 7F**; within-group, 268 ± 5 μm; across-group 273 ± 5 μm; p = 0.102, clustered Wilcoxon signed-rank test). We also compared the functional properties of pairs of neurons as a function of distance, and found that the difference in contrast modulation or engagement modulation did not depend on distance (**Figure 7G**; Pearson’s correlation with distance, difference in contrast modulation index, r = 0.003, p = 0.660; difference in engagement modulation index, r = 0.001, p = 0.960). Therefore, PPC contains neurons with diverse visuomotor response properties that are spatially intermingled.

### PPC encodes both contrast-dependent sensory signals and contrast-independent choice signals

We also analyzed the error trials to see whether stimulus and choice signals could be separately extracted from responses in PPC, and whether these signals depended on contrast (**Figure 8**). We compared False Alarm trials with Hit and Correct Reject trials, using imaging fields and sessions with at least five False Alarm trials at each contrast (7 of 11 fields, n = 391 neurons). We did not make comparisons with Miss trials given the low number of trials. The population response on FA trials was weak, but distinguishable from the response on CR trials (**Figure 8A**), especially during the choice period (**Figure 8B**; FA greater than CR for all contrasts, p < 0.05, clustered Wilcoxon signed-rank test). Using an ROC-based approach, we again found that PPC encoded both stimulus and choice with differing time courses (**Figure 8E**; stimulus selectivity significant from 1.0 to 6.6 s for all contrasts, choice selectivity significant from 2.2 to 5.6 s for all contrasts, clustered Wilcoxon signed-rank test). Interestingly, some PPC cells encoded the stimulus in a contrast-dependent manner, but also encoded the choice in a contrast-independent manner (**Figure 8C**). We quantified the contrast-dependence of the auROC index for each neuron by measuring its slope as a function of contrast (**Figure 8D**). A larger proportion of neurons exhibited significant contrast-dependence in stimulus selectivity (**Figure 8F**, 23.6% ± 19.9% of cells) compared to the proportion with significant contrast-dependence in choice selectivity (2.4% ± 1.6%). PPC may therefore simultaneously encode both contrast-dependent sensory signals and contrast-independent motor signals in the same population of neurons.

**Figure 8.**
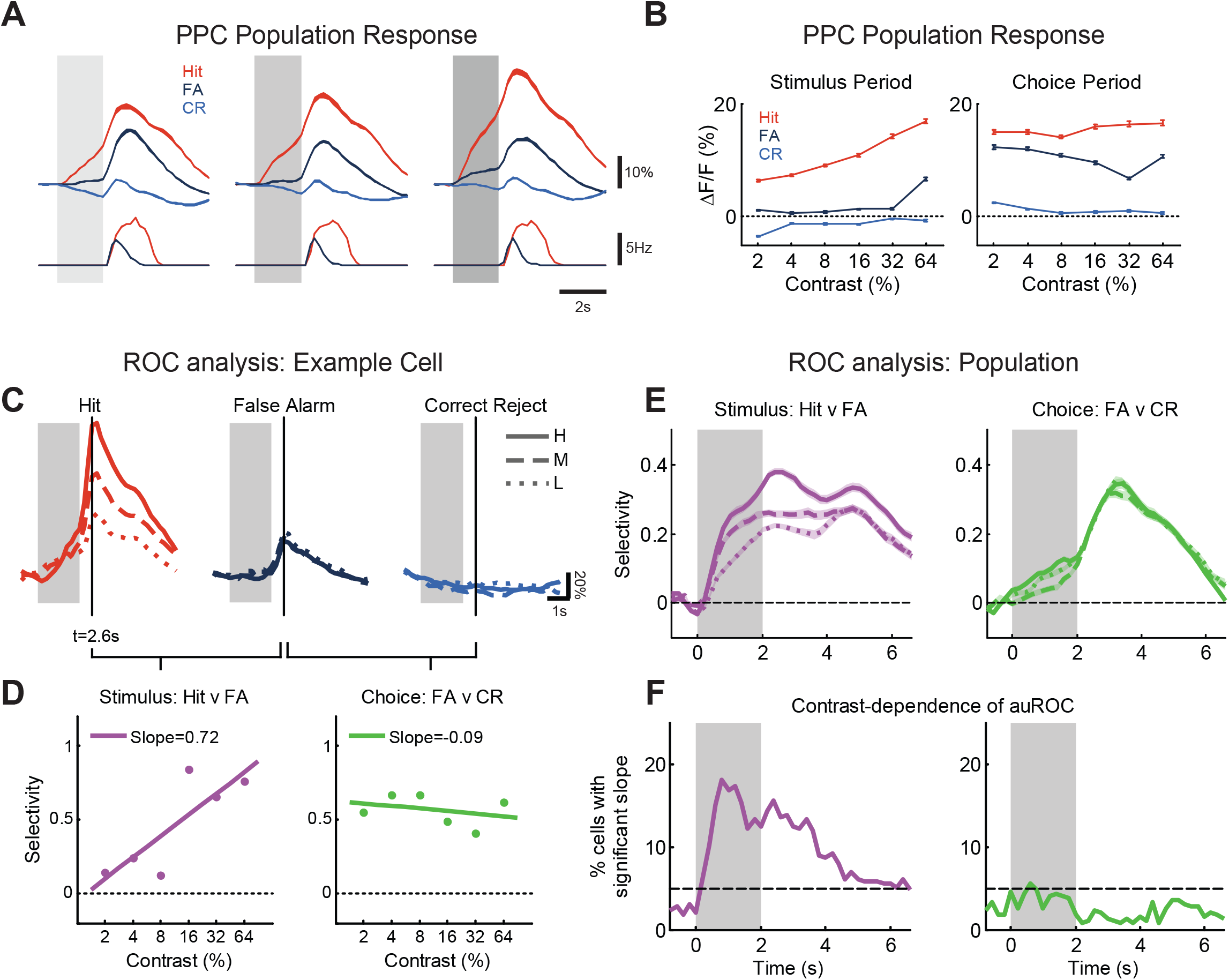
PPC encodes both contrast-dependent sensory signals and contrast-independent choice signals. (**A**) Population average of PPC responses across contrasts during Hit, False Alarm (FA), and Correct Reject (CR) trials (top). Responses are averaged across low (2 or 4%, left), medium (8 or 16%, middle), and high (32 or 64%, right) contrast. Average lick rate (bottom). Light gray shaded regions demarcate duration of stimulus. (**B**) Time-averaged population response as a function of contrast during the Stimulus period (left, 0 to 2 s) or during the Choice period (right, 2 to 3.5 s) for Hit, FA, and CR trials. Choice period responses were measured relative to preceding Stimulus period response. (**C**) Response of an example PPC neuron during Hit trials (left), False Alarm trials (middle), and Correct Reject trials (right), across different contrasts, from low (L, dotted) to medium (M, dashed), to high (H, solid). (**D**) False Alarm (FA) trials were compared with Hit trials to assess Stimulus selectivity (left), and with CR trials to assess Choice selectivity (right) at a single time-point but across multiple contrasts. Best-fit line as a function of log contrast is plotted along with its slope (arbitrary units). Stimulus selectivity is contrast-dependent, as seen with the significant positive slope, whereas choice selectivity is not. (**E**) Average stimulus selectivity (left) and choice selectivity (right) across all PPC neurons as a function of time and of contrast, from low (L, dotted) to medium (M, dashed), to high (H, solid). Shading indicates SEM. (**F**) Fraction of PPC neurons with significant contrast-dependence in stimulus selectivity (left) or choice selectivity (right), as measured by slope of auROC with respect to log contrast. Dashed line indicates fraction expected by chance.

### PPC reflects changes in stimulus-reward contingency

Thus far we have seen that PPC has a strong bias towards target stimuli during the task. Most neurons reflect the behavioral state, and a subset of neurons appear to encode the sensory stimulus in a contrast-dependent manner. Error analyses indicate that PPC neurons may encode both stimulus and choice signals, but errors can reflect other factors such as impulsivity or inattention. To more conclusively determine whether PPC neurons reflect sensory stimuli versus sensorimotor transformation, we manipulated the stimulus-reward structure of the task. By retraining mice on a reversed stimulus-reward contingency, and measuring activity from the same neurons before and after reversal, we could test whether neurons were more sensitive to stimulus identity or the learned task contingencies.

After imaging the responses of neurons in V1 and PPC in the original go/no-go task, we reversed the reward contingencies of the stimuli (**Figure 9A**). Licking in response to the original non-target stimulus (Stimulus B) was now rewarded with water, whereas licking to Stimulus A was punished with quinine. Three mice successfully learned the task after 7-11 days of training, although performance was slightly worse than before (**Figure 9B**; d-prime, original, 2.96 ± 0.50; reversed, 1.51 ± 0.22). We then measured responses from the same populations of neurons in V1 and in PPC under the reversed reward contingency, using a semi-automated procedure (Huber et al., 2012) to identify the same neurons across different imaging sessions (see **Materials and Methods** for details).

**Figure 9.**
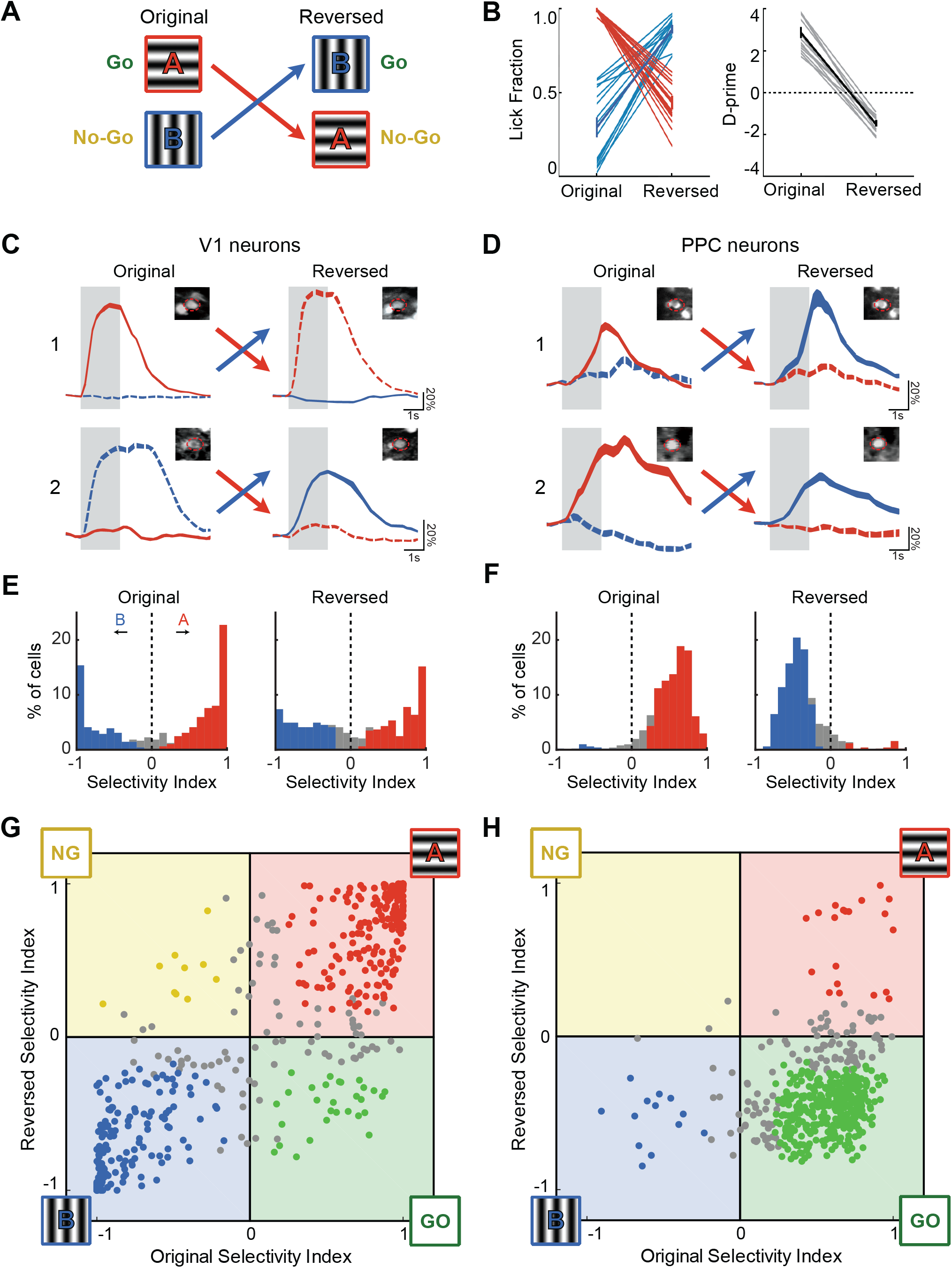
PPC reflects changes in stimulus-reward contingency. (**A**) Mice were trained on a go/no-go discrimination task with a reversed reward contingency. (**B**) Behavioral performance before and after reversal. Left, response rate for Stimulus A (red) and Stimulus B (blue). Right, d-prime assuming Stimulus A as target. Performance after reversal was significantly different from zero (p < 0.001, Wilcoxon signed-rank test). (**C**) Neural response of two example V1 neurons before and after reversal of reward contingency. Colors indicate stimulus (red, A; blue, B) and line styles indicate whether the stimulus is target (solid) or non-target (dashed). Top row, response of a neuron selective to Stimulus A both before (left) and after (right) reversal. Bottom row, response of a neuron selective to Stimulus B. Insets indicate the average projection image before and after reversal, and ROI (dashed red line) that was used to extract the neural response. (**D**) Same as (**C**) but for two PPC neurons. These neurons show a switch in selectivity with reversal of reward contingency. (**E**) Histogram of selectivity index before and after reversal for V1 neurons with significant responses both before and after reversal (n=488). Positive values indicate selectivity to the original Target, Stimulus A (red). Colored bars indicated neurons with significant selectivity. (**F**) Same as (**E**) but for PPC neurons (n=509). (**G**) Scatter plot comparing selectivity index in V1 before and after reversal. Colored points indicate neurons with significant selectivity both before and after reversal. Neurons in the first and third quadrants are stimulus-selective, as they prefer Stimulus A (red) or Stimulus B (blue) both before and after reversal. Neurons in the second and third quadrants are choice-selective, as they prefer either the Go stimulus (green) or the No-Go stimulus (yellow) in both conditions. (**H**) Same as (**G**), but for PPC.

We analyzed the selectivity of individual neurons that had significant responses both before and after reversal (V1, n = 488 cells, 8 fields in 3 mice; PPC, n= 509 cells, 8 fields in 3 mice). Many neurons in V1 that were selective to a particular stimulus remained selective to the same stimulus after reversal, whether Stimulus A or Stimulus B (**Figure 9C**). By contrast, many PPC neurons exhibited a switch in selectivity when the contingency was reversed. These neurons that were selective to target stimulus A during the original task became selective to the new target stimulus B after reversal (**Figure 9D**).

We first examined the overall selectivity in neurons in V1 and PPC (**Figure 9E, F**). For V1 neurons, the selectivity index did not change significantly after reversal (before, 0.161 ± 0.034; after, 0.095 ± 0.030; p = 0.256, clustered Wilcoxon signed-rank test), with a relatively symmetrical distribution of neurons selective to either stimulus A or B. However, in PPC, selectivity was dramatically altered with reversal of reward contingency (before, 0.513 ± 0.013; after, -0.373 ± 0.013; p = 4.15 × 10^-3^, clustered Wilcoxon signed-rank test), with the majority of responsive neurons preferring the new target stimulus B after reversal.

We then plotted the selectivity of individual neurons before and after reversal against each other as a scatter plot (**Figure 9G, H**). Purely stimulus-selective neurons will remain close to the unity line and in the first (bottom-left) and third (top-right) quadrants, whereas neurons that are sensitive to the reward contingency will lie in either the second quadrant (top-left; for no-go selective cells) or the fourth quadrant (bottom-right; for go-selective cells). V1 neurons were strongly stimulus-selective, with the majority (74.3% ± 5.2% of cells, **Figure 10A**, top) of neurons lying within the first and third quadrants (**Figure 9G**). By contrast, the majority (66.8% ± 13.0% of cells, **Figure 10A**, bottom) of PPC neurons were found in the fourth quadrant, indicating a preference for the rewarded target stimulus, regardless of its actual identity (**Figure 9H**). These results indicate that the majority of PPC neurons indeed reflect the learned task contingencies.

**Figure 10.**
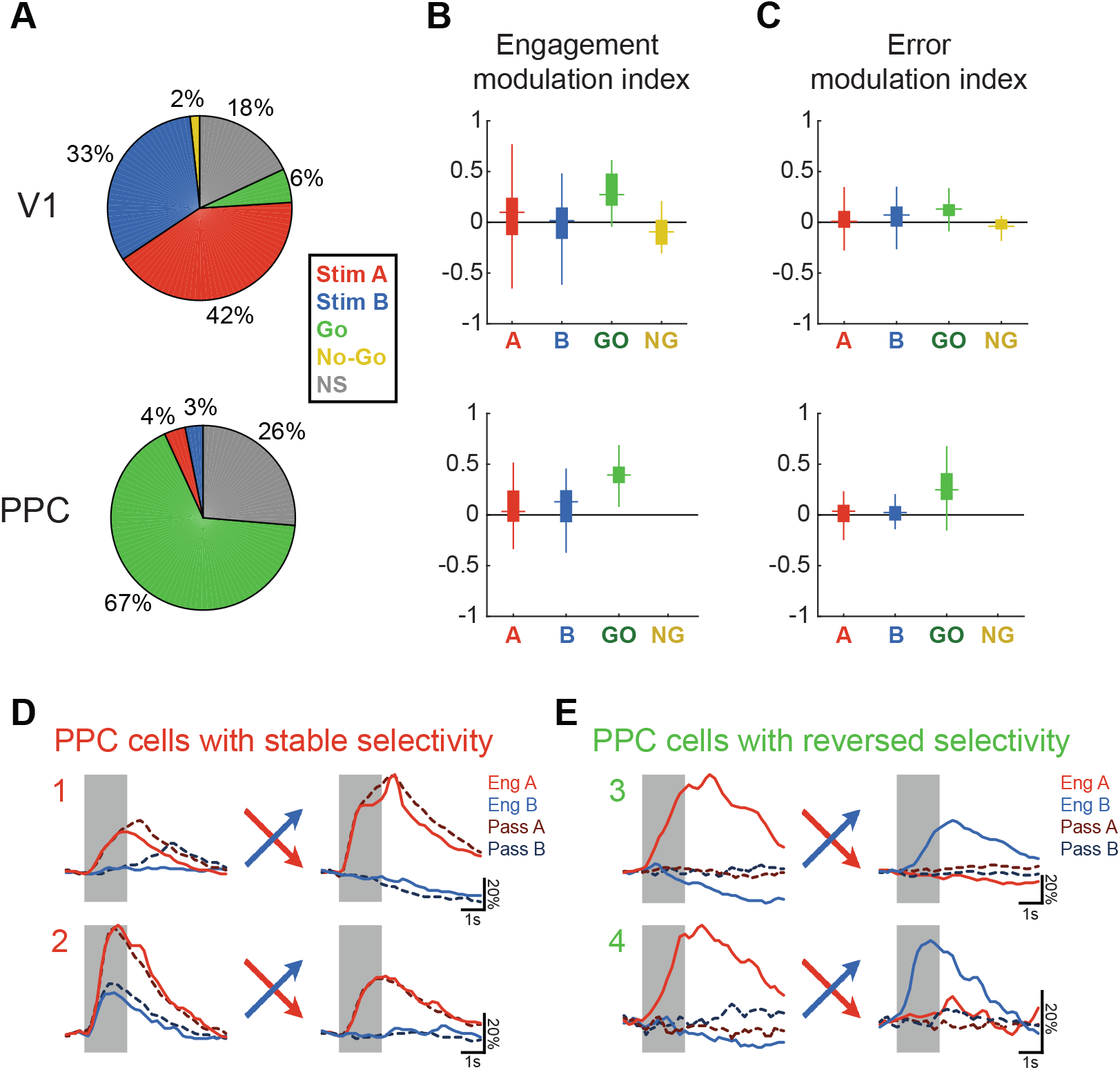
PPC cells with reversed selectivity exhibit stronger modulation by engagement and by error trials. (**A**) Proportion of significantly responsive V1 (top) and PPC (bottom) neurons with stable stimulus selectivity (Stim A, 0°; or Stim B, 90°) or reversed selectivity (Go or No-Go). Cells with nonsignificant selectivity either before or after reversal are categorized as non-selective (NS). (**B**) Boxplots (showing median, interquartile range, and range) of engagement modulation index for each response group, computed as the mean before and after reversal. “Go”-selective neurons (green) show stronger engagement modulation, i.e. weaker passive responses, than stimulus-selective neurons (red or blue). (**C**) Boxplots of error modulation index for each response group, computed by comparing False Alarm trials with Correct Reject trials both before and after reversal. “Go”-selective PPC neurons show stronger error modulation (green) than stimulus-selective neurons (red or blue). (**D**) Responses of two PPC neurons with stable selectivity to Stimulus A both before and after reversal of reward contingency. Colors indicate stimulus (red, A; blue, B) and dark, dashed lines indicate passive trials. These neurons show a robust passive response in both conditions, and have a low engagement modulation index. (**E**) Same as (**D**) but for two PPC neurons with reversed selectivity after reversal of reward contingency. These neurons show virtually no passive response in both conditions, and have a high engagement modulation index.

### PPC cells with reversed selectivity exhibit stronger modulation by engagement and by error trials

A small subset of PPC neurons (6.9% ± 3.7%) did show stable stimulus selectivity before and after reversal. Given that a minority of neurons also exhibited stimulus-driven passive responses that were also insensitive to behavioral choice on error trials, we wondered whether these different properties (dynamic selectivity, engagement modulation, and error modulation) were related and could be used to separate functional cell types. In other words, if PPC neurons could indeed be separated into groups of “sensory” and “motor” neurons, then stimulus selectivity after reversal should be predictive of modulation strength with engagement and choice.

To test this hypothesis, we measured the strength of engagement modulation in both stimulus-selective (to either Stimulus A or B) neurons and go-selective neurons, as defined by their selectivity before and after reversal (**Figure 10A**). We found that go-selective PPC neurons had higher (near significant) modulation by engagement than stimulus-selective neurons (go-selective, 0.393 ± 0.007; stimulus-selective, 0.084 ± 0.040; p = 0.065, clustered Wilcoxon rank-sum test; **Figure 10B**, bottom). Indeed, many PPC neurons that maintained selectivity to Stimulus A after reversal also had strong passive responses (**Figure 10D**), and many neurons with reversed selectivity were strongly gated by engagement (**Figure 10E**). We also tested to see whether a PPC neuron’s responses on error trials related to its selectivity after reversal. Go-selective PPC neurons had significantly higher error modulation (computed by comparing False Alarm trials to Correct Reject trials) compared to stimulus-selective neurons (go-selective, 0.273 ± 0.009; stimulus-selective, 0.019 ± 0.019; p = 0.040, clustered Wilcoxon rank-sum test; **Figure 10C**). Interestingly, the small subset of go-selective neurons in V1 (5.9% ± 2.1% of cells) also differed from stimulus-selective V1 neurons in their degree of engagement (**Figure 10B**, top, p = 0.016) and error (**Figure 10C**, top, p = 0.021) modulation, suggesting that these different properties are strongly related and can be used to distinguish functional cell types across different areas.

These findings demonstrate distinct subsets of neurons within PPC. One subset of neurons faithfully reflects the sensory input, both in passive conditions and after learning a new reward contingency. The larger proportion of PPC neurons, however, switch selectivity after retraining, and are strongly modulated by behavioral state. The flexibility and heterogeneity of PPC responses suggests a role for PPC in the mapping of sensory inputs onto appropriate motor actions.

## Discussion

We developed a head-fixed visual decision task for mice with separate stimulus and response epochs, and used population imaging to investigate the role of PPC in perceptual decisions. Our key findings are that PPC encodes both sensory and motor signals across a heterogeneous pool of neurons, and that its activity can change drastically depending on task performance and demands. Together these results suggest that mouse PPC is responsible for neither pure sensory processing, nor for the control of motor output, but rather is important for the decision process itself – the process of mapping sensation to action.

### Mouse PPC is more than an extrastriate visual area

The small size of the mouse brain has made it difficult to identify precise borders between different functional areas. PPC in the rodent, as classically defined by its thalamic inputs (Reep et al., 1994), is located in the region between the more posterior V1 and the more anterior somatosensory cortex. However, this location is essentially where both anatomical (Wang and Burkhalter, 2007; Wang et al., 2012) and functional (Marshel et al., 2011; Garrett et al., 2014) mapping studies have identified the retinotopically-organized areas RL, A, and AM. The degree to which PPC overlaps with these secondary visual areas is a matter of debate. Some have argued that rodent PPC may have more in common with primate extrastriate cortex, given that inactivation specifically disrupts sensitivity on visual decisions (Licata et al., 2017). Indeed, the stereotaxic coordinates used by us (Goard et al., 2016) and others (Harvey et al., 2012; Morcos and Harvey, 2016) to target mouse PPC most directly overlap with area AM, which exhibits directionally-tuned responses even under anesthesia (Marshel et al., 2011). But is PPC simply a sensory visual area?

We have presented multiple pieces of evidence that point to a role for PPC beyond mere sensory processing. First, activity in PPC is strongly dependent on behavioral state, with only a minority of task-responsive PPC neurons exhibiting significant responses during passive viewing of stimuli (Figure 2F). Secondly, the selectivity of PPC neurons is strongly biased toward target stimuli (Figure 2E). This bias is likely due to the asymmetry of the Go/No-go paradigm, and may reflect a learned association of stimulus and reward (Fitzgerald et al., 2013). Third, PPC responses are modulated during error trials (Figure 3, Figure 8). Information about the eventual choice of the animal can be decoded from the activity of PPC, as previously shown in both mice (Harvey et al., 2012; Goard et al., 2016) and rats (Raposo et al., 2014). Finally, and most conclusively, the biased selectivity of most PPC neurons towards target stimuli is dramatically reversed when the animal is retrained on a different reward contingency (Figure 9). Together these results demonstrate that the stimulus-period responses in PPC are task-dependent and not purely sensory.

### PPC activity does not merely reflect movement

One may argue that the activity patterns we observed in PPC can be most parsimoniously explained as movement or action-planning related signals. After all, movement-related activity would be present only during engagement, it would exhibit choice selectivity, and it would change with reward contingency. It is unlikely that PPC is directly involved in executing motor plans, as we have previously shown that optogenetic inactivation of PPC during the response period, or even during the delay between stimulus and response, has no effect on behavior (Goard et al., 2016). However, the possibility remains that the PPC responses recorded in our task reflect planning- or movement-related signals that originate elsewhere. Although some PPC neurons (29%) do show activity that appears motor-related, due to their contrast-independent nature (Figure 6E), we provide evidence that PPC is not a purely motor area either.

First, passive visual stimulation does induce a response in some PPC neurons (20%), as previously shown in parietal area AM of anesthetized mice (Marshel et al., 2011). This subset of neurons tends to stably reflect the stimulus even with changes in reward contingency (Figure 10). Second, we find a subset of PPC neurons that reflect the target stimulus on Miss trials when the animal fails to lick, even though such neurons are inactive during passive viewing (Figure 3). Finally, the responses of many PPC neurons (43%) shows modulation with stimulus contrast, even for the same decision and motor output (Figure 6E). This is reminiscent of previous primate (Shadlen and Newsome, 2001) and rodent (Hanks et al., 2015) PPC studies, where it has been shown that responses vary with the strength of incoming sensory evidence.

### Strength of noise correlations in V1 and PPC

PPC neurons can be sensitive to both sensory and motor signals, but a linear model incorporating these signals can only partially explain the variability of PPC responses (average 20% variance explained). While there are potentially many other unmeasured variables that drive this variability, one source appears to be the activity of other neurons in the network. As previously shown by others (Zohary et al., 1994; Cohen and Maunsell, 2009), a population of neurons can express strong inter-neuronal dependencies that result in correlated activity fluctuations from trial to trial, (but see Ecker et al., 2010; Renart et al., 2010). Our results extend these findings by demonstrating that noise correlations are more prominent in PPC than in V1, are reduced by task engagement, and can improve prediction performance when incorporated into linear models of responses. An intriguing possibility is that highly recurrent cortical regions such as PPC and prefrontal cortex may be more subject to noise correlations than sensory regions with strong feedforward drive.

One caveat is that these models and measured correlations were based on calcium responses, an indirect measure of spiking activity. While our lab (Rikhye and Sur, 2015) and others (Hofer et al., 2011; Cossell et al., 2015; Lur et al., 2016) have observed relative changes in correlations using calcium imaging that are qualitatively similar to those obtained from electrophysiological recordings, it must be acknowledged that the absolute strength of correlations cannot be directly related across recording modalities. Nonetheless, we observed specific relative differences between PPC and V1, and between Engaged and Passive conditions, which resulted in differing improvements in model performance.

### Heterogeneity of PPC responses

In every task condition reported here, PPC responses were heterogeneous. A subset of PPC neurons have significant passive visual responses and are modulated by contrast in both engaged and passive conditions. These neurons maintain their stimulus selectivity even after reversal of reward contingency. These “sensory” neurons are spatially intermingled with the larger proportion of “motor” neurons that have task-gated responses, weak contrast modulation, and altered selectivity after contingency reversal.

Heterogeneous response properties have been previously reported in both primate (Bennur and Gold, 2011; Meister et al., 2013) and rat (Raposo et al., 2014) PPC during decision tasks. Our work adds to this literature, and additionally provides evidence that such heterogeneous responses exist in a spatially intermingled fashion within PPC of the mouse. Do these response types form bona-fide cell classes or does PPC represent a category-free population, as others have proposed (Raposo et al., 2014)? Although we do find that various properties (such as modulation by engagement, contrast, and reversal) are correlated with one another, more work needs to be done to determine whether functional neuronal subgroups are truly separable. Future work can take advantage of the tools available for the mouse system to determine whether these functional properties correlate with molecular markers of cell identity (Kvitsiani et al., 2013; Pinto and Dan, 2015) or with projection target (Chen et al., 2013a; Li et al., 2015).

### Conclusion and outlook

Our results demonstrate that mouse PPC encodes sensory, decision, and motor variables, suggesting a role in sensorimotor transformation during perceptual decisions. Our reversal experiments reveal a strong experience-dependent plasticity of representations in PPC. Although previous studies have similarly shown that representations in PPC can be altered with re-training (Freedman and Assad, 2006), our work unequivocally shows such changes in individual PPC neurons. Taking advantage of the stability and consistency of two-photon imaging, we demonstrate that not only PPC as a whole, but also many individual neurons show a remarkable switch in selectivity after reversal of the task reward structure.

The mouse posterior parietal cortex encodes behaviorally-relevant variables in a highly task-dependent manner, in analogy to prior work in primates. Our understanding of how decisions are computed and sensorimotor transformations are made will be greatly aided by future circuit-level analyses of PPC function in this powerful model system (Carandini and Churchland, 2013).

## Acknowledgements

This work was supported by the NIH (M.J.G., F32-EY023523 and K99-MH104259; M.S., R01-EY007023 and U01-NS090473); the NSF (G.N.P., Graduate Research Fellowship; M.S., EF1451125); the Simons Center for the Social Brain (M.S.), and the Picower Institute Innovation Fund (M.J.G.; M.S.). We thank J. Sharma, A. Boesch, V. Li, C. Le, J. Krizan, V. Breton-Provencher, and T. Emery for technical assistance; R. Huda, M. Hu, H. Sugihara, and C. D. Harvey for helpful discussions and/or comments on the manuscript; L. L. Looger, J. Akerboom, D. S. Kim, and the Genetically-Encoded Neuronal Indicator and Effector (GENIE) Project at Janelia Farm Research Campus Howard Hughes Medical Institute for generating and characterizing GCaMP6 variants.

## Author contributions

G.N.P. and M.J.G. designed experiments with input from M.S.; M.J.G. performed the surgeries; M.J.G. and G.N.P. performed the imaging experiments; M.J.G., G.N.P., J.W., and B.C. performed the behavioral training; G.N.P. developed the behavior software; G.N.P. analyzed the data with comments from M.J.G. and M.S.; G.N.P., M.J.G., and M.S. wrote the manuscript.

